# Dchs2, a novel Fat4 ligand, is required for photoreceptor organization and outer limiting membrane integrity in the mouse retina

**DOI:** 10.64898/2026.08.05.742812

**Authors:** Jennysue Kasiah, Didier Hodzic, Nicole Liscio, Helen McNeill

## Abstract

Fat and Dachsous cadherins are large transmembrane proteins that regulate tissue growth and planar cell polarity across species. In mammals, Fat4 is known to bind Dchs1, but whether it also binds Dchs2 is unclear. Here we show that Dchs2 binds Fat4 in a trans-heterophilic manner at cell-cell contacts, mirroring the Fat4-Dchs1 interaction. In the developing mouse retina, Fat4 and Dchs1 expression peaks embryonically and declines after birth, whereas Dchs2 expression begins postnatally and persists into adulthood, with all three co-expressed from birth through the second postnatal week. While *Fat4* and *Dchs1* mutants show no defects in retinal lamination or cell-type composition, loss of *Dchs2* increases number of cones, reduces Müller glia number, and results in disorganization of photoreceptor outer segments. Electron microscopy shows that *Dchs2* mutant photoreceptors fail to form the regular nuclear columns seen in controls, with disrupted stacking and breaks in the outer limiting membrane. These findings identify Dchs2 as a novel Fat4 ligand and reveal a distinct, non-redundant requirement for *Dchs2* in photoreceptor organization and outer limiting membrane integrity in the mature retina.

## Introduction

Cadherins are adhesion molecules that modulate tissue morphogenesis via cell sorting and migration (Angst et al., 2001; Hulpiau and van Roy, 2009) to sustain proper development. The Fat/Dachsous cadherin superfamily are enormous, evolutionarily conserved adhesion molecules involved in tissue growth and patterning (Bryant et al., 1988; Clark et al., 1995; Fulford and McNeill, 2020; Mao et al., 2011; Tanoue and Takeichi, 2005). *Drosophila* Fat (Ft) regulates tissue growth via the Hippo pathway (Bennett and Harvey, 2006; Bosch et al., 2014; Cho et al., 2006; Fulford et al., 2023; Silva et al., 2006; Willecke et al., 2006), planar cell polarity (PCP) via a non-canonical Ft-Ds pathway (Ambegaonkar et al., 2012; Bosveld et al., 2012; Brittle et al., 2022; Matakatsu and Blair, 2004, 2006, 2012; Matis and Axelrod, 2013; Sharma and McNeill, 2013; Zhao et al., 2013), and mitochondrial function via Complex I (Sing et al., 2014). *Drosophila* Dachsous (Ds) also regulates tissue growth (Baena-Lopez et al., 2008; Willecke et al., 2008) and PCP (Ambegaonkar et al., 2012; Bosveld et al., 2012; Brittle et al., 2022; Matakatsu and Blair, 2004, 2006, 2012). A key characteristic of these proteins is that they act as receptor-ligand pairs through trans-heterophilic bonds across cells. (Clark et al., 1995; Ma et al., 2003; Matakatsu and Blair, 2004).

Mammals express four Fat cadherins (*Fat1-4*) and two Dachsous (*Dchs1* and *Dchs2*) cadherins. *Fat4*, the ortholog of *ft*, is a large, single-pass transmembrane protein (Fig. 1A) that regulates tissue growth in various cancers (Abuderman et al., 2020; Cai et al., 2015; Che et al., 2019; Du et al., 2013; Hou et al., 2016; Jiang et al., 2018; Li et al., 2020; Li et al., 2019; Ma et al., 2016; Malgundkar et al., 2020; Mao et al., 2022; Pan et al., 2020; Sun et al., 2018; Wei et al., 2019), as well as a variety of developmental processes including skeletal system formation (Crespo-Enriquez et al., 2019; Mao et al., 2016), kidney development (Bagherie-Lachidan et al., 2015; Saburi et al., 2008; Zhang et al., 2019), neuronal migration (Badouel et al., 2015; Zakaria et al., 2014), and cerebral cortex apical membrane organization (Ishiuchi et al., 2009). *Dchs1* and *Dchs2* are orthologs of *ds*, consisting of large, single-pass transmembrane proteins (Fig. 1A) that share ∼30% intracellular domain sequence similarity to each other (Rock et al., 2005). Dchs1 and Fat4 interact trans-heterophilically at cell-cell contacts (Loza et al., 2017; Medina et al., 2023; Strutt et al., 2024).

**Figure 1.**
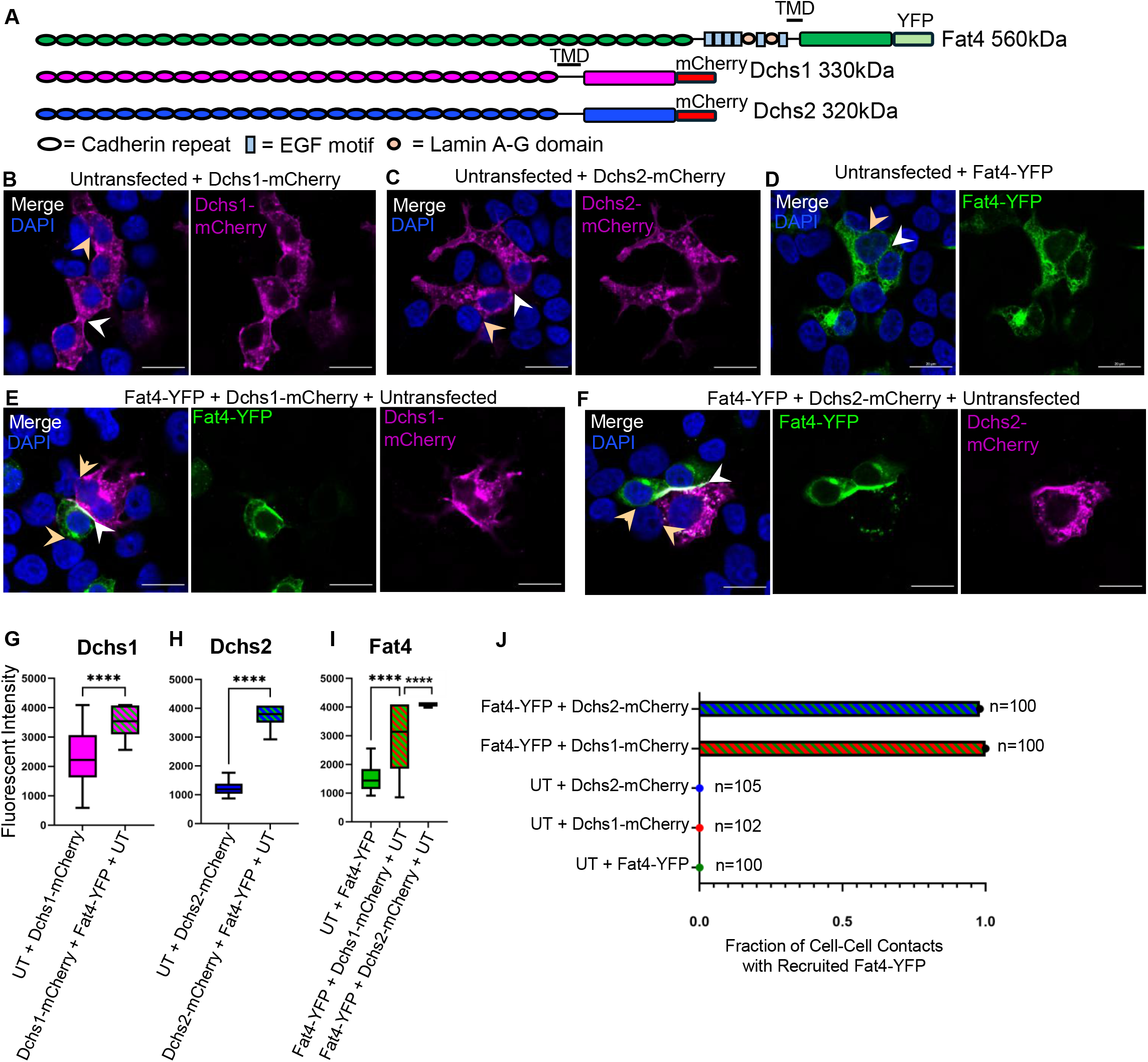
Dchs2 binds Fat4 trans-heterophically. **A.** Schematic representation of Fat4, Dchs1, Dchs2 cadherins with location of either YFP or mCherry, and molecular weights. TMD= transmembrane domain **B-D**. Untransfected (UT) HEK293 cells were co-cultured with Dchs1-mCherry, Dchs2-mCherry or Fat4-YFP expressing HEK293 cells and analyzed for protein accumulation at cell-cell contacts. No protein accumulation was observed with Dchs1-mCherry (n=102), Dchs2-mCherry (n=105) or Fat4-YFP (n=100). n=total number of cells counted across 3 replicates. Tan arrows indicate contacts between Fat4, Dchs1, or Dchs2 expressing cells and UT cells. White arrows indicate contacts were two transfected cells touch **E.** Fat4-YFP expressing cells were co-cultured with Dchs1-mCherry and UT cells and analyzed for protein accumulation at cell-cell contacts out of n=100 contacts, 100 showed protein accumulation measured by fluorescent intensity. **F.** Fat4-YFP expressing cells were co-cultured with Dchs2-mCherry and UT cells and analyzed for protein accumulation at cell-cell contacts. Out of n=100 contacts, 98 showed protein accumulation measured by fluorescent intensity. **G-I.** Fluorescent intensity was quantitated by drawing a line that bisects both cells crossing through the cell-cell contact of each contact imaged. Fluorescent intensity across the 100 contacts counted per condition were averaged and plotted. **J.** Fraction of cell-cell contacts with recruited Fat4-YFP. Scale bars=20µM. Results presented are mean fluorescent intensities across cell-cell contacts with error bars representing SEM. ****p<0.001

Unlike *Fat4*^KO^ or *Dchs1*^KO^ mice, *Dchs2*^KO^ mice are viable and fertile (Bagherie-Lachidan et al., 2015). *Dchs2*^KO^ mice have pituitary hypoplasia and infundibular defects (Lodge et al., 2020). Notably, loss of both *Dchs1* and *Dchs2* has a synergistic effect in restricting the nephron progenitor pool in the mouse kidney, suggesting some redundancy (Bagherie-Lachidan et al., 2015), and loss of Dchs1/2 is also synergistic in zebrafish (Castelvecchi et al., 2026). Loss of *Dchs1/*2 also results in the mislocalization of Vangl1, causing altered positioning of the basal body in node cells (Sai et al., 2022). Whether Dchs2 acts through a homophilic or heterophilic interaction is unknown, and its downstream pathway is not well understood.

Fat cadherins regulate many aspects of eye development. In the lens epithelium, polarity and proliferation of lens progenitor cells are maintained via *Fat1* (Sugiyama et al., 2015), and proper amacrine cell migration and cone-bipolar cell synapse connection via *Fat3* (Aviles et al., 2022; Aviles et al., 2025; Krol et al., 2016). *Fat4* expression has been characterized in the developing eye via *in situ* hybridization from embryonic day 9.5 through 18.5. *Fat4* is first detected at E10.5 in lens pit cells, and by E12.5 *Fat4* expression is confined to the anterior portion of the lens vesicle. Additionally, at E12.5 *Fat4* is seen in the optic cup. At E14.5 *Fat4* is localized at the distal margin of the optic cup. By E18.5 *Fat4* marks a distinct lamina in the inner portion of the neural retina. Loss of *Fat4* was also examined in the embryonic eye up to E18.5, however, no defects were reported in either eye formation or lens development (Sugiyama et al., 2015).

Here we show that Dchs2 acts as a ligand for Fat4 *in vitro*: cell mixing assays reveal that Fat4 and Dchs2 accumulate at cell-cell contacts in a trans-heterophilic fashion, whereas Dchs2 does not bind homophilically. Fat4, Dchs1 and Dchs2 proteins are present in the developing mouse retina, with distinct temporal expression dynamics. Fat4 and Dchs1expression are highest embryonically and decline after birth. In contrast, Dchs2 expression is low at birth and increases during retinal maturation. Dchs1, Dchs2 and Fat4 are co-expressed from post-natal day 0-15 (P0-P15). Loss of *Fat4* or *Dchs1* led to decreased apical Par3, with no overall alterations in apical-basal cell polarity. Histological and immunofluorescent examination of adult *Fat4*, *Dchs1* and *Dchs2* KO retinas revealed no overt morphological phenotypes, or dramatic changes in cell fate, although *Dchs2* mutants had a mild increase in cone photoreceptors, and a decrease of Müller glial cells. Strikingly, loss of *Dchs2* specifically disrupted photoreceptor inner segment organization and the columnar stacking of photoreceptor nuclei and compromised the outer limiting membrane (OLM). Loss of *Fat4* or *Dchs1* did not affect this organization. Taken together, these data reveal a specific role for *Dchs2* in photoreceptor organization.

## Results

### Fat4 and Dchs2 interact at cell-cell contacts

Fat4 and Dchs1 act as binding partners *in vivo* in the embryonic cerebral cortex (Cappello et al., 2013) and *in vitro* in HEK 293T cells (Loza et al., 2017) and L cells (Ishiuchi et al., 2009). To determine if Dchs2 can act as a ligand for Fat4, we generated stable HEK293 cell lines expressing either full-length Fat4-YFP, Dchs1-mCherry or Dchs2-mCherry (Fig.1B, C, D). Consistent with previous reports that Fat4 can recruit Dchs1 to cell contacts, Fat4-YFP accumulated at cell contacts with Dchs1-mCherry expressing cells (Fig. 1E, G, J) but not at contacts with untransfected HEK293 cells. Similarly, cells expressing Dchs2-mCherry showed no accumulation at cell contacts with other Dchs2-mCherry expressing cells, nor at contacts with untransfected HEK293 cells (Fig 1C) indicating Dchs2 does not bind homophilically. Importantly, co-culture of Fat4-YFP and Dchs2-mCherry expressing cells revealed that both proteins accumulate at cell-cell contacts (Figure 1F, H, J), indicating that Dchs2 can bind and recruit Fat4 to cell-cell contacts. These data indicate that Dchs2 can function as a binding partner for Fat4.

### Fat4, Dchs1 and Dchs2 are expressed in the developing retina

To examine the developmental expression pattern of Fat4 (∼560kDa) in the mouse retina, retinal lysates from E14.5, P0, P5, P15, and P30 wild-type eyes and or from a control P0 *Fat4^KO^* mice were analyzed by western blot with a Fat4 antibody detecting the intracellular domain of Fat4 (Fig 2E). As shown in Figure 2A-A’, Fat4 expression peaks at E14.5 but becomes extremely reduced in the adult retina.

**Figure 2.**
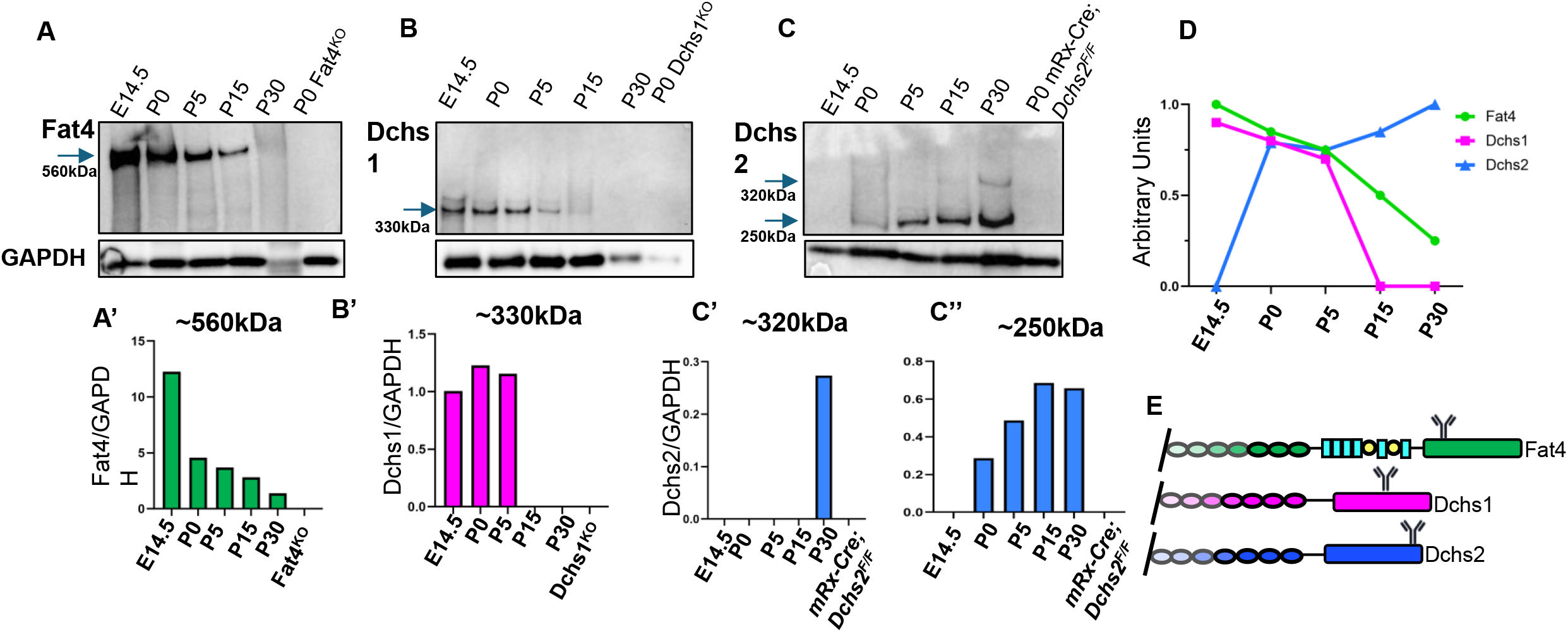
Fat4 and Dchs1 show highest expression embryonically while Dchs2 expression is highest postnatally. **A-C.** Time course western blots wildtype P0, P5, P15, P30 and a P0 Fat4^KO^ (A), Dchs1^KO^ (B) or mRx-Cre; Dchs2^F/F^ (C) control. **A’-C’.** Quantitation of westerns blots showing a gradual decrease of Fat4 as the retina matures, a decrease of Dchs1 starting at P5, and an increase in Dchs2 as the retina matures starting at P0. **D.** Simplified diagram depicting the trends of Fat4, Dchs1, and Dchs2 during retina development. **E.** Schematic of where each antibody detects either Fat4, Dchs1 or Dchs2.

The same wild type lysates – as well as a lysate prepared from P0 Dchs1 null retinas – were further examined for the expression of endogenous Dchs1 (∼330 kDa) using an antibody directed towards the C-terminus of Dchs1. Interestingly, Dchs1 and Fat4 temporal expression patterns were similar, with a gradual decrease after birth (Fig. 2A-2B’, J).

To probe for endogenous Dchs2 (∼320 kDa), we generated a Dchs2 antibody recognizing the C-terminal region (see materials and methods). Western blot analysis revealed that Dchs2 expression is not detectable at E14.5 but gradually increases postnatally with a prominent ∼250kDa band starting at P0 and a weaker band of ∼320kDa starting at P15 (Fig 2C-2C”). This data suggests that either a splice variant or cleaved product (∼250kDa) is the predominant Dchs2 form from P0-15 and that the predicted full-length Dchs2 (∼320kDa) is not present until the retina fully matures at P30. Taken together these data highlight the dynamic expression patterns of endogenous Fat4, Dchs1 and Dchs2 during eye development, with Fat4 and Dchs1 highest expression in early retina development, and Dchs2 highest expression in the mature retina, with overlap of all three proteins from P0-P15 (Fig. 2D). Given that we have shown that Fat4 can bind Dchs2, these data suggest that Fat4 may change binding partners during eye development.

### Fat4 localizes to the RPE, neuroblast and lens epithelium

Previous studies examining the localization of Fat4 in the embryonic eye revealed that Fat4 localizes apically in embryonic epithelial tissues at adherens junctions (Ishiuchi et al., 2009). We examined E14.5 wildtype eyes for apical localization of Fat4. Using a commercially available Fat4 antibody that detects the intracellular domain (Fig. 2E), we were able to visualize Fat4 in three structures within the eye: the retinal pigment epithelium (RPE), the neuroblast layer, and the lens epithelium (Fig. 3A-E). Interestingly, the RPE and neuroblast are two structures with interacting apical surfaces; here we see a strong Fat4 signal, validated by the loss of signal in the *Fat4^KO^* eye (Fig. 3B). Finer examination of the RPE and neuroblast interfaces revealed that Fat4 localizes to both apical surfaces (Fig. 3C, E”). Fat4 signal in the lens epithelium, however, is enriched at lateral membranes, with no apical enrichment (Fig. 3 D-E’), a unique pattern for Fat4.

**Figure 3.**
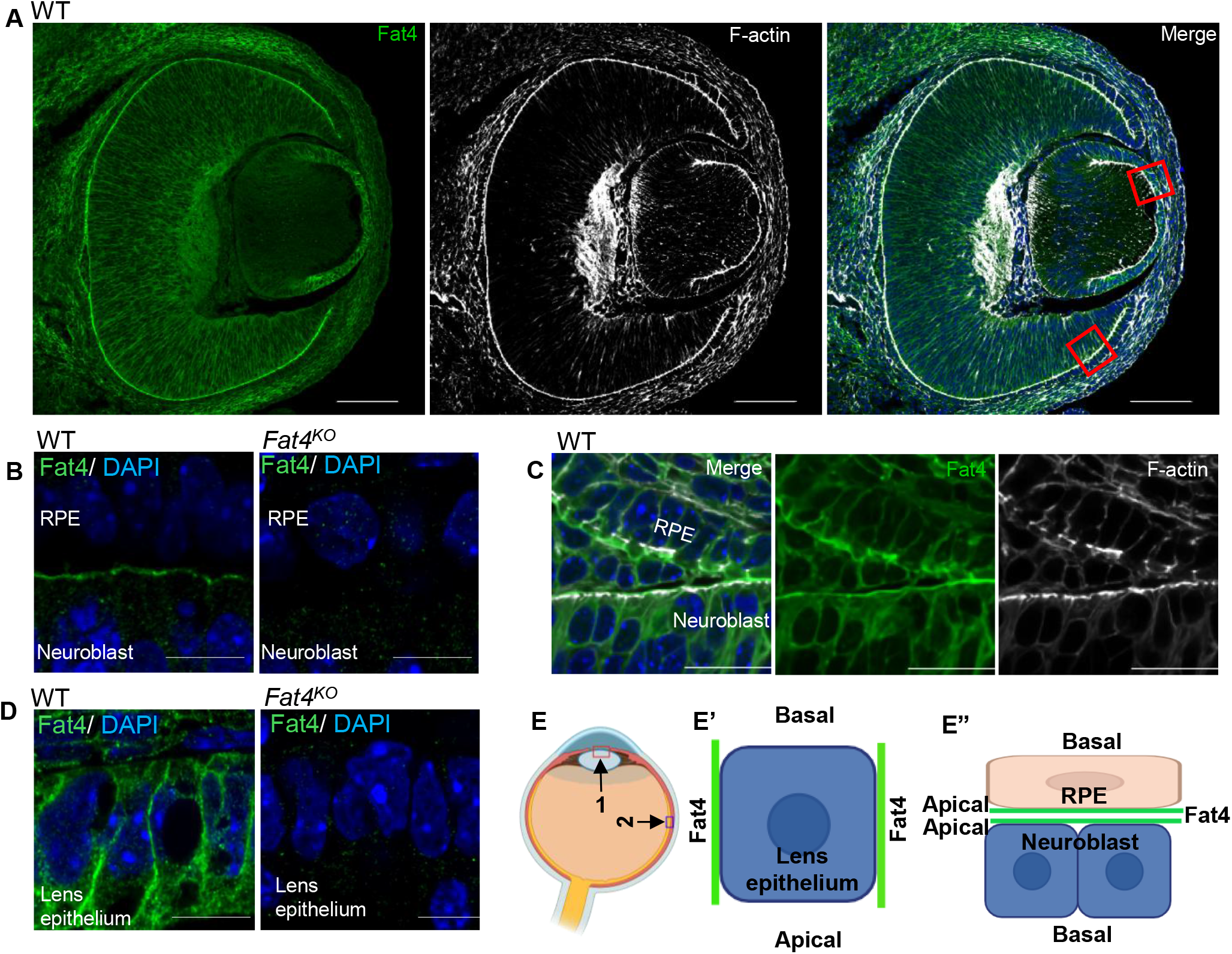
Fat4 localizes to the RPE/neuroblast and lens epithelium. **A.** E14.5 wildtype eye labeled for Fat4 (green) and F-actin (white). Scale bar = 100µm **B.** Zoomed in image of the RPE/neuroblast in wild type (WT) and Fat4^KO^ retinas labeled for Fat4 (green) and nuclei (blue). Scale bar = 20µm **C.** A zoomed in image of the RPE and neuroblast showing that Fat4 (green) localizes to both the apical surface of the RPE and the apical surface of the neuroblast. Apical surface the RPE and neuroblast are marked with F-actin (white). Scale bar = 20µm. **D.** Zoomed in image of the lens epithelium showing that Fat4 (green) localizes laterally. Scale bar = 20µm. **E.** Schematic showing the location of the lens epithelium (arrow 1) or the RPE/neuroblast (arrow 2). **E’.** Schematic showing lateral Fat4 localization of a lens epithelial cell. **E”.** Schematic of the RPE/Neuroblast apical interface showing Fat4 localization at both apical surfaces.

### Loss of *Fat4* does not affect Yap/Taz localization or ciliary body growth in the retina

A core characteristic of *Drosophila ft* is its ability to regulate tissue growth and size via the Hippo pathway (Baena-Lopez et al., 2008; Bennett and Harvey, 2006; Bosch et al., 2014; Cho et al., 2006; Matakatsu and Blair, 2012; Silva et al., 2006; Willecke et al., 2006). However, in mammals *Fat4’*s connection to the Hippo pathway is tissue and timing context dependent (Che et al., 2019; Du et al., 2013; Kuta et al., 2016; Malgundkar et al., 2020). Previous work demonstrated that the mammalian retinal pigment epithelium (RPE) and ciliary body size are controlled by *Nf2,* an up-stream regulator of the Hippo pathway (Moon et al., 2018). We asked if *Fat4* could be modulating the hippo pathway in the RPE/ciliary body. Using P30 mRx-Cre; *Fat4^F^*^/F^ and *Fat4^F^*^/F^ animals we assessed readouts for altered Hippo pathway function: nuclear Yap/Taz localization (Supp 1A-B) and growth of the ciliary body (Supp 1C-D). There was no significant difference in nuclear Yap/Taz between control and mRx-Cre; *Fat4^F/F^* animals (Supp. 1B). We did not detect any overgrowth of the ciliary body at P30. These data indicate that neither the RPE nor ciliary body need *Fat4* for Hippo pathway regulation.

### Fat4 and Par3 bind *in vitro* and co-localize to the apical neuroepithelium

Previous work from our lab using Fat4-FLAG expressing HEK293 cells and AP-MS analysis identified PARD3 (Par3) as a putative Fat4-interacting protein (Badouel et al., 2015). To confirm the binding of Fat4 and Par3 we generated FLAG-Fat4-ICD constructs and conducted FLAG co-immunoprecipitation (coIP) assays. Using lysates generated from the coIP’s, we assessed binding of Par3 to Fat4 constructs via western blot. We found that Par3 strongly binds to the full Fat4 ICD (FLAG-Fat4-ICD) but only weakly to constructs with C-terminal deletions (Supp. 2A-B). These data confirm that Fat4 interacts with Par3 through the C-terminal region of the Fat4 cytoplasmic domain. We next asked if Par3 and Fat4 colocalize to the apical surface of the retina neuroepithelium. Because both Par3 and Fat4 antibodies are produced in rabbit, we generated an endogenous, fluorescently tagged *Fat4* allele containing a tension sensor of mTFP1, a spider silk linker and Venus, referred to henceforth as *Fat4^Venus^*, to co-label both proteins in the retina (Fig. 4A). Fat4^Venus^ homozygous animals are viable and healthy, with no detectable defects. We first compared the localization pattern of Fat4^Venus^, using an anti-GFP antibody, to Fat4 antibody staining and found consistent apical localization of the Venus signal in the RPE/neuroblast and lateral signals in the lens epithelium (Fig 4B-D), confirming the usefulness of the Fat4^Venus^ animals in reflecting endogenous Fat4 distribution. E14.5 *Fat4^Venus^* eyes were co-stained for Fat4 and Par3, revealing colocalization at the apical surface of the neuroepithelium (Fig. 4E).

**Figure 4.**
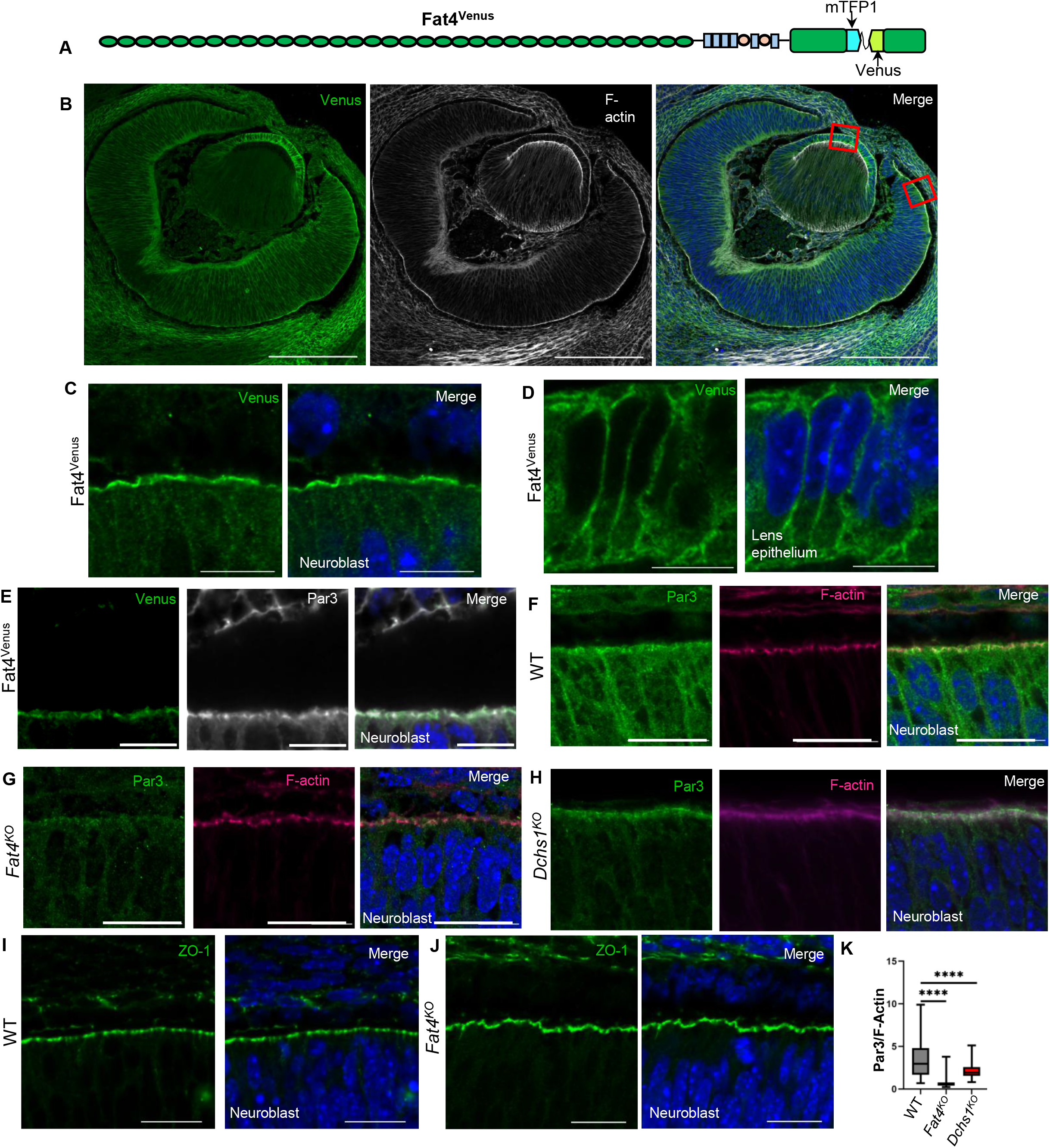
Loss of *Fat4 or Dchs1* reduces apical Par3 but does not affect apico-basal polarity. **A.** Schematic of the *Fat4^Venus^* allele showing the insertion of the tension sensor cassette into the intracellular domain of Fat4. **B.** Whole E14.5 eye labeled with anti-GFP (Venus) and Phalloidin (F-actin). Red boxes on the merged image indicate the area, RPE/neuroblast or lens, that are highlighted in panels **C** and **D,** respectively. **C.** Fat4^Venus^ signal localizes to the RPE/ neuroblast. **D.** Fat4^Venus^ localizes laterally in the lens epithelium. **E.** E14.5 Fat4^Venus^ retina labeled with Venus (green) and Par3 (white), showing colocalization of Fat4 and Par3 at the apical surface of the neuroblast. **F-H.** WT, *Fat4^KO^* or *Dchs1^KO^* E14.5 retinas labeled for Par3 (green) and F-actin (Pink). **I-J.** E14.5 WT and *Fat4^KO^* retinas labeled with ZO-1 (green) showing no change in localization of ZO-1 in *Fat4^KO^* conditions **K.** Quantitation of **F-H** assessing apical intensity of Par3 reveals a significant decrease in apical Par3 in mutant conditions, with loss of *Fat4* causing the greatest reduction of Par3. ****= p<0.0001.

### Loss of *Fat4* or *Dchs1* reduced apical Par3 in the embryonic retina

In the vertebrate eye, Par3 localizes to the apical surface of retina neuroepithelium to organize apico-basal polarity and lamination (Kanda et al., 2013). We asked if loss of *Fat4* alters the localization of Par3 in the embryonic retina. Examination of E14.5 wildtype and *Fat4^KO^* eyes for Par3 and F-actin localization revealed a significant decrease in Par3 at the apical surface of the neuroepithelium, but no changes to F-actin localization, MCT1, ZO-1 or P cadherin localization or overall cell morphology (Fig. 4F-L and Supp. 2C-D). These data suggest that Fat4 helps recruit or stabilize Par3 at the apical surface, but that loss of Fat4 is not sufficient to cause disruptions to apical-basal polarity in the retina. Because Fat4 can bind Dchs1, we also examined E14.5 retinas of *Dchs1* null animals for changes in Par3 localization, we found similar reduction in Par3 fluorescence intensity in *Dchs1*^KO^ retinas (Fig 4H-K), with loss of *Fat4* showing the greatest reduction of apical Par3 (Fig. 4J).

### Loss of *Fat4* does not affect retina morphology or cell type composition

We next asked if retina morphology was impacted by the loss of *Fat4*. Because *Fat4^KO^* animals die shortly after birth, we generated *Fat4* conditional knockouts using the mRx-Cre system to examine mature retinas. mRx-Cre turns on at E8 and targets the retina, the RPE and portions of the forebrain (Klimova et al., 2013). To assess retinal morphology, six thickness measurements were taken: total retinal thickness (Retina), outer nuclear layer (ONL), inner nuclear layer (INL), outer plexiform layer (OPL), inner plexiform layer (IPL), and inner/outer photoreceptor segments (segments). Examining P30 mRx-Cre; *Fat4*^F/F^ histological eye sections, we found no change in the thickness of retinal layers (Fig 5A-C). We also examined whether the relative populations of retinal cell types were affected. We stained the eyes for five different cell types: cones (cone arrestin), rods (rhodopsin), rod bipolar cells (PKC_α_), horizontal cells (calbindin D28K) and Müller Glia (glutamine synthetase). We detected no change in cell composition in P30 mRx-Cre; *Fat4*^F/F^ eyes (Fig. 5D-D’). These data indicate that *Fat4* is not necessary for proper retinal lamination or cell-type composition.

**Figure 5.**
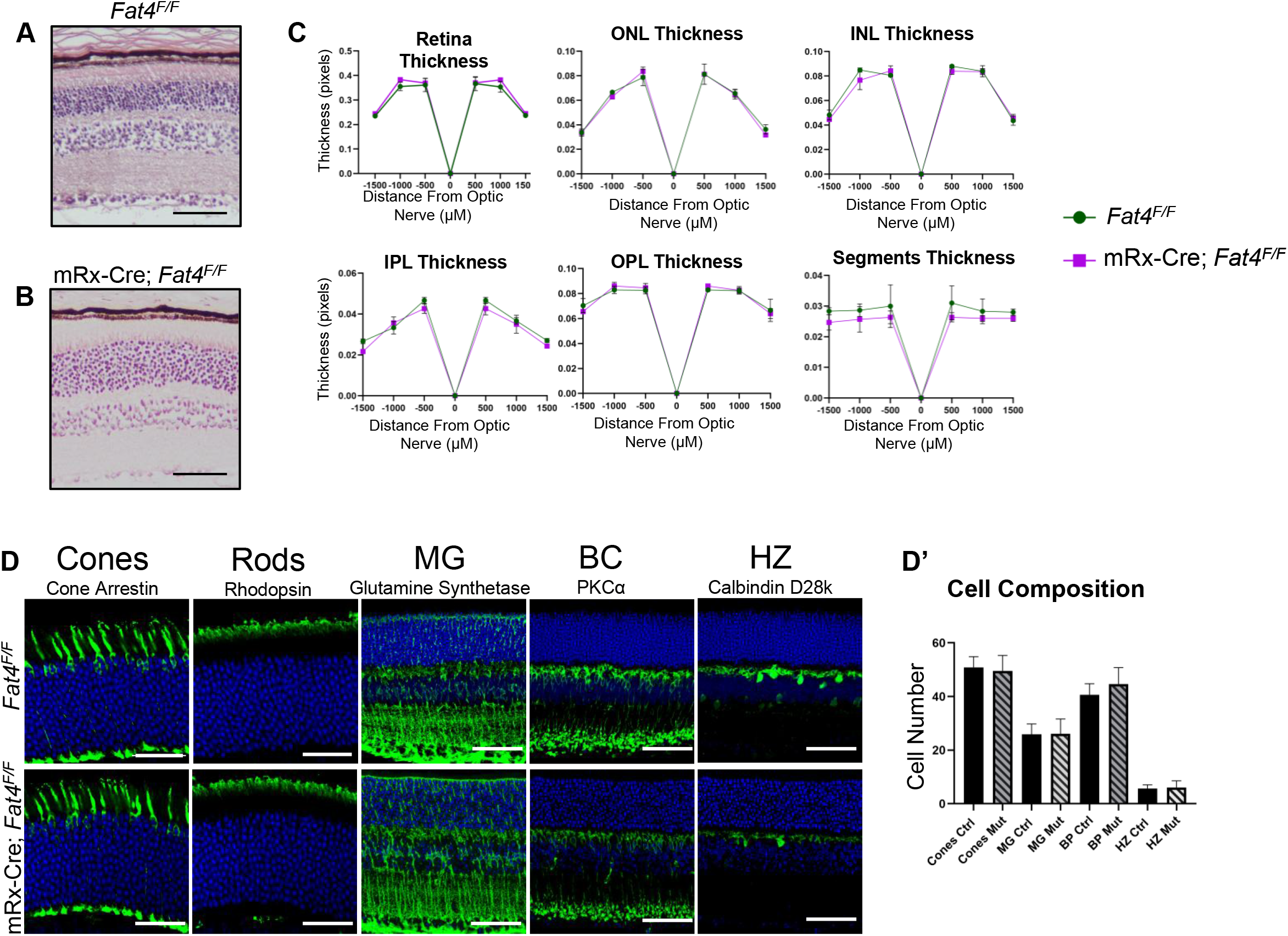
Loss of Fat4 does not affect retina morphology or cell-type composition. **A.** P30 H&E of *Fat4^F/F^* retina Scale bar = 50µm. **B.** P30 H&E of mRx-Cre; *Fat4^F/F^* retina. Scale bar = 50µm **C.** Quantitation of the layers of the retina showing no significant changes between *Fat4^F/F^* (green) and mRx-Cre; *Fat4^F/F^*(purple). **D.** Fluorescent images of P30 retinas labeling cones, rods, Müller glia (MG), bipolar cells (BC) and horizontal cells (HZ). Scale bars = 20µm **D’.** quantitation of number of the different cell types between *Fat4^F/F^* and mRx-Cre; *Fat4^F/F^*.

### *Dchs1* loss does not affect retina morphology or cell type composition

As shown in Figure 2, Dchs1 is highly expressed in the embryonic retina. However, limitations of available antibodies made it difficult to determine its retinal localization. We therefore turned to a Dchs1-HA mouse (Byerly et al., 2025) and determined that Dchs1 localizes to the apical surface of the retina neuroepithelium, similar to the expression pattern of Fat4 (Fig. 6A-B’).

**Figure 6.**
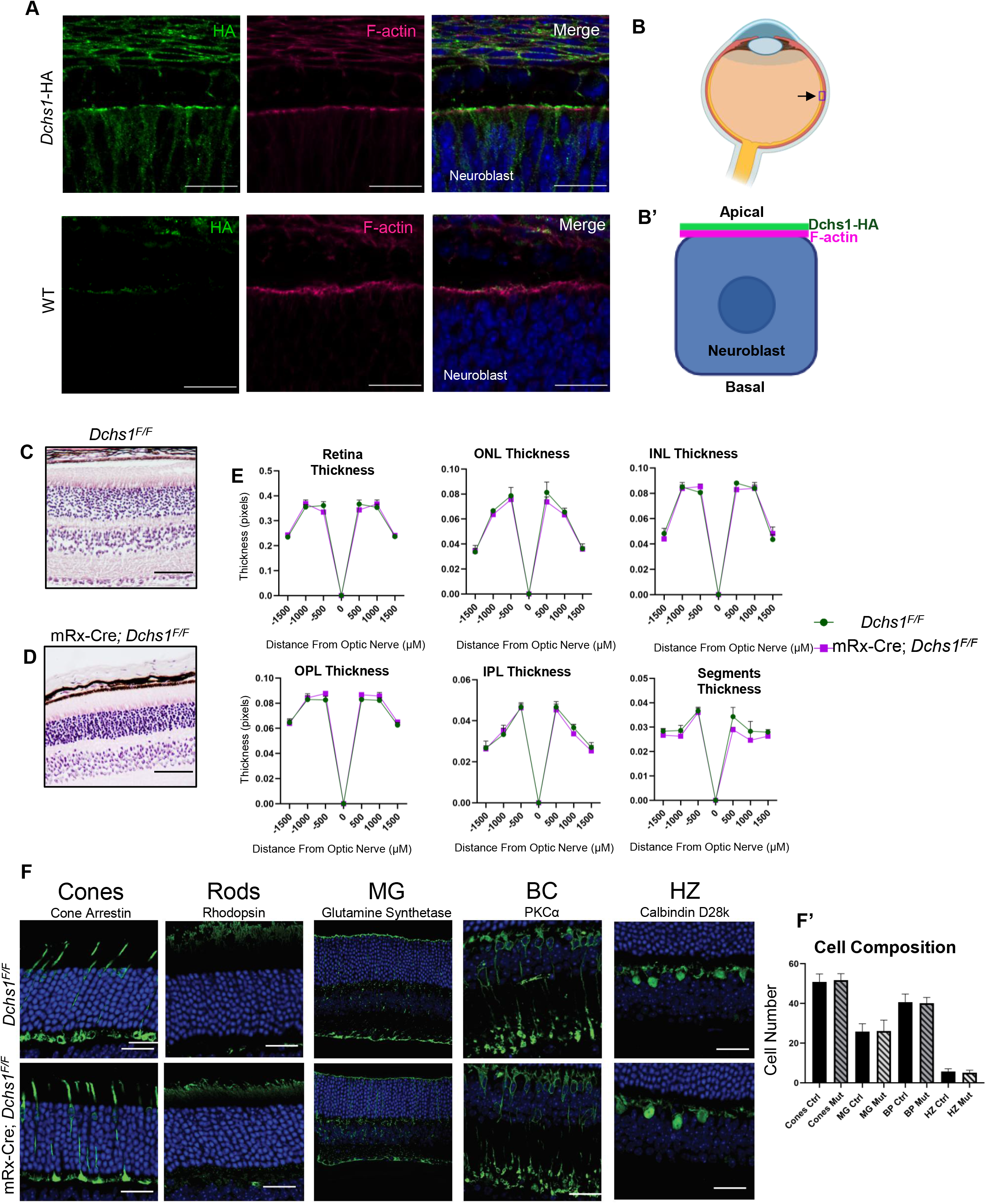
Dchs1 localizes to the neuroblast and RPE but loss of *Dchs1* does not affect retina morphology or cell-type composition. **A.** Zoomed in image of *Dchs1*-HA and WT neuroblast (HA green) (F-actin pink). Scale bar = 20µm. **B.** Schematic of the eye showing the location of the retina where Dchs1 localizes **B’.** Schematic of a neuroepithelial cell showing Dchs1 (green) localizing apically above F-actin (pink). **C.** P30 H&E of *Dchs1^F/F^* retina Scale bar = 50µm. **D.** P30 H&E of mRx-Cre; *Dchs1^F/F^* retina. Scale bar = 50µm **E.** Quantitation of the layers of the retina showing no significant changes between *Dchs1^F/F^* (green) and mRx-Cre; *Dchs1^F/F^*(purple). **F.** Fluorescent images labeling P30 retinas for cones, rods, Müller glia (MG), bipolar cells (BC) and horizontal cells (HZ). **F’.** Quantitation of number of the different cell types between *Dchs1^F/F^* and mRx-Cre; *Dchs1^F/F^*.

We next asked if *Dchs1* regulates retina morphology or cell-type development. Similar to *Fat4*, loss of *Dchs1* is perinatal lethal (Mao et al., 2011), so we generated conditional knockouts (mRx-cre; *Dchs1*^F/F^), and assessed the thickness of the retina, ONL, INL, OPL, IPL and segments at P30. Analysis of these layers revealed no significant changes in mRx-cre; *Dchs1*^F/F^ eyes compared to *Dchs1*^F/F^ (Fig 6B-D).

Further analysis of cones, rods, horizontal cells, bipolar cells, and Müller glia revealed no significant changes between mRx-cre; *Dchs1*^F/F^ and *Dchs1*^F/F^ (Fig. 6D-D’). Taken together, these data indicate that *Dchs1* is expressed in the embryonic retina but is non-essential for retinal morphology or cell type composition.

### Loss of *Dchs2* mildly affects cone and Müller glia populations but does not affect retina lamination

While Dchs2 can be detected in the postnatal retina by Western blot (Fig. 2), we were unable to determine the subcellular localization of Dchs2 due to lack of specific antibodies. Using mRx-Cre; *Dchs2*^F/F^ animals, we found no change to gross retina morphology of mutants compared to controls via thickness assessment of the retina, ONL, INL, OPL, IPL or outer/inner segments (Fig. 7A-C). Cell type composition analysis, however, showed a significant increase in the number of cones in mRx-Cre; *Dchs2*^F/F^ animals compared to *Dchs2^F/F^* controls (Fig 7D-D’). We also observed a significant decrease in the number of Müller glia (Fig. 7D’).

**Figure 7.**
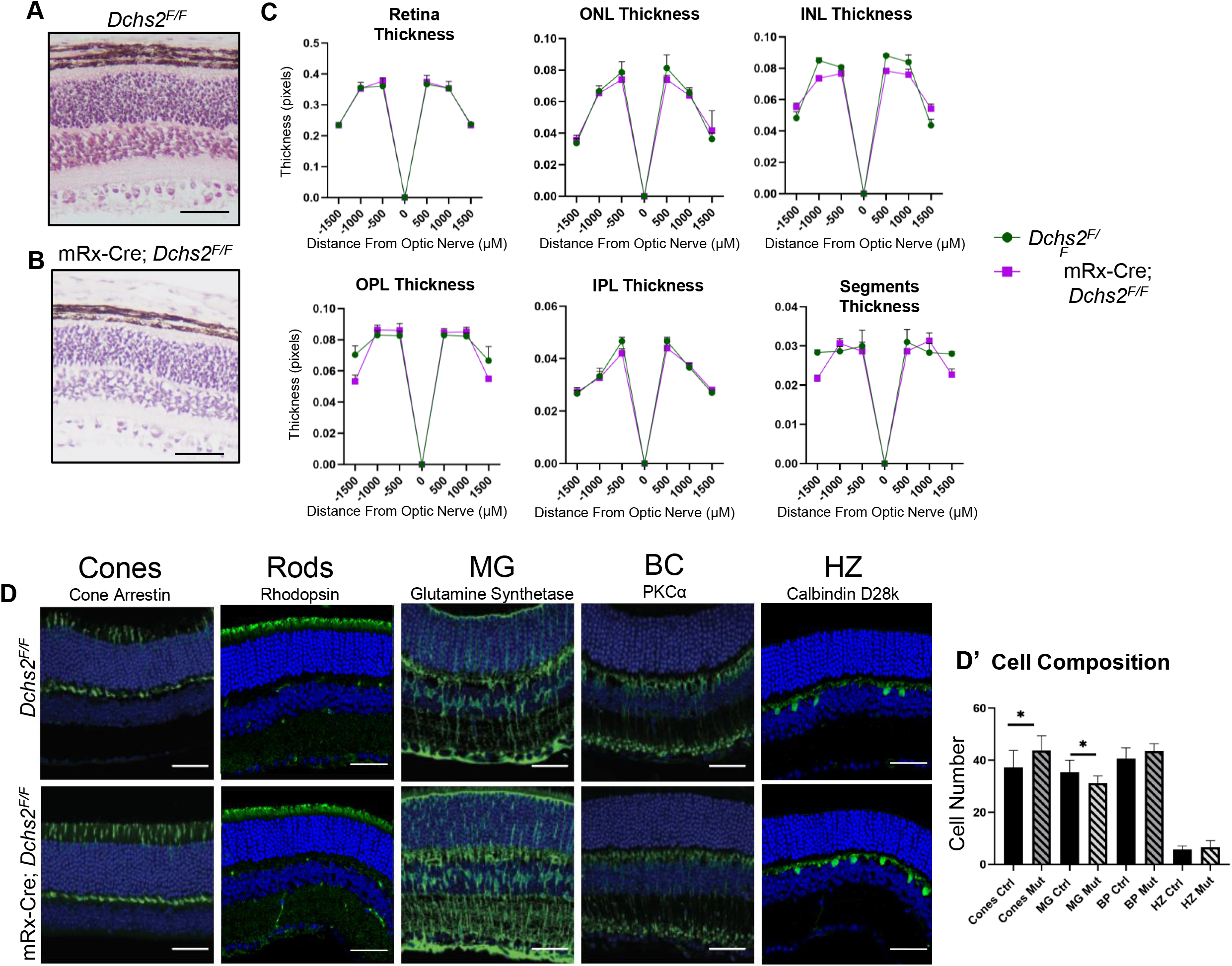
Loss of Dchs2 does not affect retina morphology but results in a decrease of cones and Müller glia. **A.** P30 H&E of *Dchs2^F/F^* retina Scale bar = 50µm. **B.** P30 H&E of mRx-Cre; *Dchs2^F/F^* retina. Scale bar = 50µm **C.** Quantitation of the layers of the retina showing no significant changes between *Fat4^F/F^* (green) and mRx-Cre; *Dchs2^F/F^*(purple). **D.** Fluorescent images labeling P30 retinas for cones, rods, Müller glia (MG), bipolar cells (BC) and horizontal cells (HZ). **D’.** quantitation of number of the different cell types between *Dchs2^F/F^* and mRx-Cre; *Dchs2^F/F^.* Cones * = p 0.0369. MG * = p 0.0298

### Loss of Dchs2 results in disorganization of photoreceptors

Close examination of P30 mRx-Cre; *Dchs2*^F/F^ retinas suggested ONL defects. To assess the ONL/photoreceptor segments we stained with antibodies against glutamine synthetase, which marks Müller glia projections, and retinoschisin (RS1), a protein needed for retina structural integrity which marks the photoreceptor inner segments (Molday et al., 2007).This analysis revealed that mRx-Cre; *Dchs2*^F/F^ mutant eyes have disorganized photoreceptor inner segments and weaker Müller glia staining at the outer limiting membrane (OLM) (Fig. 8A). The OLM is an adherens junction belt that connects photoreceptors to Müller glia, which is necessary to maintain proper retina morphology (Tworig and Feller, 2021). In addition to weaker glutamine synthetase staining, we observed breaks in F-actin and ZO-1 in the mRx-Cre; *Dchs2*^F/F^ eyes that were associated with nuclei pushing into the OLM (Fig. 8B). These data indicate that the OLM is compromised in animals that lack *Dchs2*. Additionally, we examined P30 mRx-Cre; *Dchs1*^F/F^ and mRx-Cre; *Fat4*^F/F^ eyes and found no changes to glutamine synthetase, F-actin or RS1 (Supp. 3A-B).

**Figure 8.**
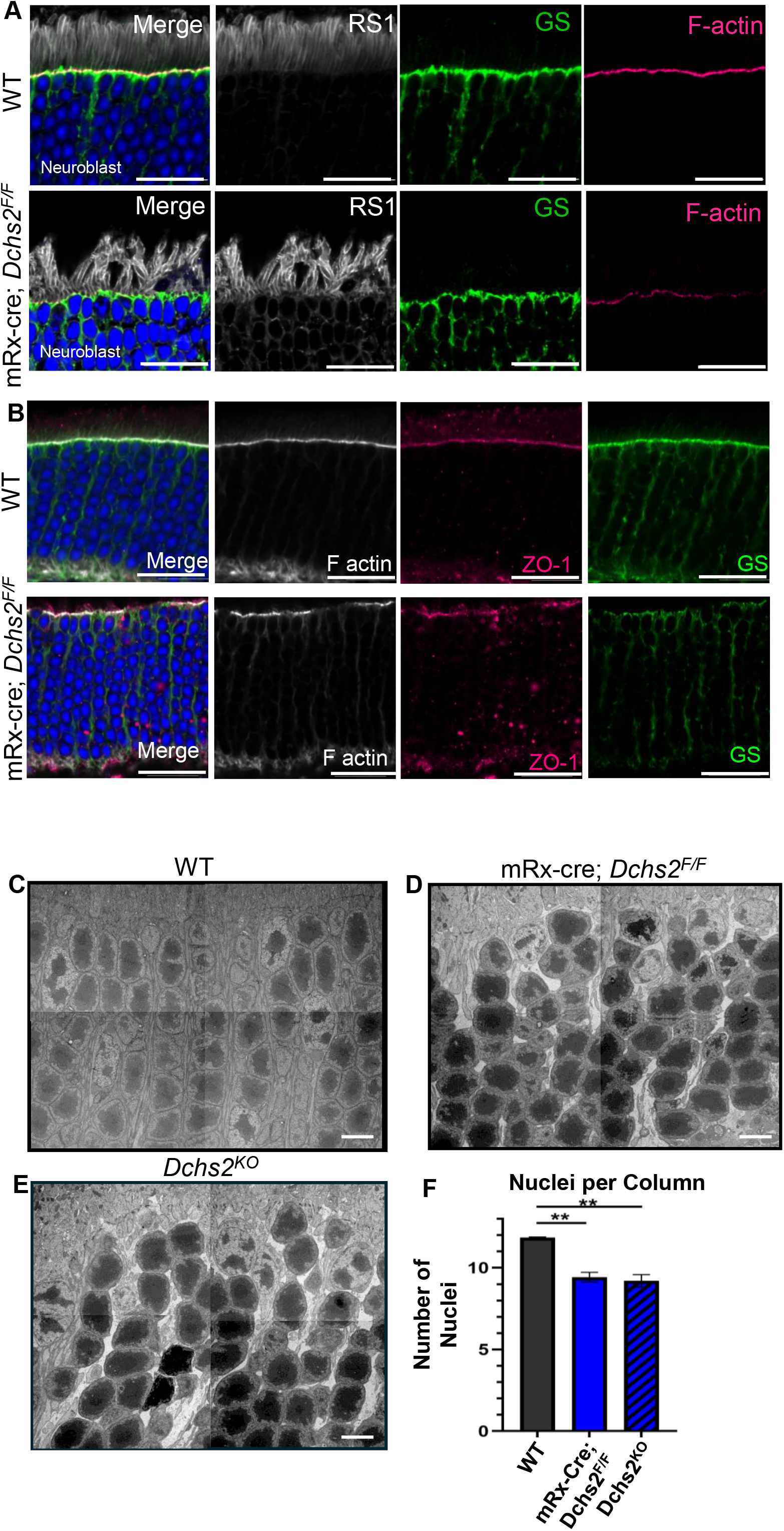
Loss of *Dchs2* disrupts organization of photoreceptor inner segments and columnar structure. **A.** P30 WT and mRx-Cre; *Dchs2^F/F^* retinas labeled with RS1 in white (inner segments), glutamine synthetase (GS) in green (Müller glia projections), and F-actin in pink. Photoreceptor inner segments of mRx-Cre; *Dchs2^F/F^* retinas show significant changes in organization compared to WT controls. **B.** WT and mRx-Cre; *Dchs2^F/F^* retinas labeled with GS (green), F-actin (white), and ZO-1 (pink). mRx-Cre; *Dchs2^F/F^* retinas have nuclei that break through the outer limiting membrane (OLM), causing disruptions in the F-actin and ZO-1. **C-E.** TEM images of WT **(C.)**, mRx-Cre; *Dchs2^F/F^* **(D.)** and Dchs2^KO^ **(E.)** retinas focusing on the columnar structure of the. Scale bars = 2µm. **F.** Quantitation of nuclei per column showing a significant decrease in the number of nuclei in mRx-Cre; *Dchs2^F/F^* (**** = p 0.0046) and in *Dchs2^KO^* (**** = p 0.0064) retinas.

Further assessment of P30 mRx-Cre; *Dchs2*^F/F^ mutants using electron microscopy (EM) showed substantial changes in photoreceptor nuclei organization as well as disruptions to the ONL. In wildtype, the ONL is organized into columns that contain 12-13 nuclei stacked on one another. We found that mRx-Cre; *Dchs2*^F/F^ mutant retinas do not form these stacked columns; rather, the nuclei are arranged haphazardly, forming columns with fewer nuclei per column compared to the controls (Fig. 8C-F). *Dchs2^KO^* animals are fertile and viable (Bagherie-Lachidan et al., 2015). Therefore, we also assessed retinas from these animals and found similar disorganization to the retinal structure, indicating that these effects are due to loss of *Dchs2* and not off target Cre effects (Fig. 8D-F).

## Discussion

Ft and Ds act as receptor and ligand for signaling in *Drosophila*. In mammals, there are two Ds homologs, *Dchs1* and *Dchs2*. Fat4 and Dchs1 are known to interact as receptor-ligand and are needed in a variety of developmental processes including the skeletal system (Bagherie-Lachidan et al., 2015; Crespo-Enriquez et al., 2019; Kuta et al., 2016; Mao et al., 2016; Mao et al., 2011), heart (Byerly et al., 2025; Mao et al., 2011; Ragni et al., 2017), kidneys (Bagherie-Lachidan et al., 2015; Mao et al., 2011; Saburi et al., 2008; Zhang et al., 2019) and brain (Badouel et al., 2015; Ishiuchi et al., 2009; Zakaria et al., 2014). However, the ligand for Dchs2 was previously unknown. In this study, we generated full length cDNAs for Dchs2 and demonstrated that Dchs2 can act as a binding partner for Fat4. Using an *in vitro* system, we show that Dchs2 does not interact with itself in a homophilic manner; rather it acts in a trans-heterophilic fashion to bind Fat4, similar to the trans-heterophilic binding of Fat4 and Dchs1. It is unknown how the binding of Dchs2 to Fat4 affects downstream signaling of *Fat4*, and whether *Dchs2* plays a role in the Fat/Dchs PCP pathway. Further investigation is needed to explore the molecular effects *Dchs2* may have on *Fat4* signaling. Whether Dchs2 also binds other Fat family members (Fat1, Fat2, Fat3), several of which are expressed in the eye, remains to be tested.

We also identified distinct temporal expression patterns for Fat4, Dchs1 and Dchs2 during retinal development. Fat4 and Dchs1 levels are highest embryonically and decline after birth. Interestingly, Dchs2 is low at birth and rises during retinal maturation. Given that there is a temporal switch from Dchs1 to Dchs2 as the retina matures, it is possible that the ligand for Fat4 changes from Dchs1 during development to Dchs2 during early postnatal retina maturation. It is unknown how the binding of Fat4 to Dchs2 affects downstream pathways of *Fat4*, however our data indicate that at least some of the functions of *Dchs2* are *Fat4*-independent.

Using immunostaining and analysis of mice with an *in vivo* knock-in Venus tag in *Fat4*, we found that Fat4 localizes to the apical surface of both the RPE and retina neuroepithelium, similar to the embryonic cerebral cortex. Interestingly, in the lens epithelium Fat4 is not enriched apically but rather localizes laterally. This is a unique pattern for Fat4, and further exploration is needed to understand the role Fat4 plays in lens development and maturation. Fat4 in the lens poses many exciting avenues for exploration as organization of the lens epithelium and orientation of the lens fibers is coordinated via the core PCP pathway and loss of the Wnt/Fz antagonist *Sfrp2* results in improper alignment of the lens fibers (Adler and Belecky-Adams, 2002; Sugiyama et al., 2011; Sugiyama et al., 2015; Sugiyama et al., 2010). It is intriguing to speculate that lens PCP could be regulated by a Fat/Dchs PCP pathway.

Previous work from our lab using proteomic screens identified Par3 as an interactor of Fat4 (Badouel et al., 2015). Here we confirmed with co-immunoprecipitation that Fat4 binds Par3 *in vitro* and co-localizes with Par3 at the apical surface of the neuroepithelium. Furthermore, loss of *Fat4* reduces apical Par3, although it does not impact overall apical-basal organization of the neuroepithelium. The fact that loss of Fat4 does not abolish Par3 localization implies other Par3 interactors may also recruit and/or stabilize Par3 such as aPKC (Izumi et al., 1998; Nagai-Tamai et al., 2002), Par6 (Joberty et al., 2000; Lin et al., 2000), and junction adhesion molecules (JAM) (Ebnet et al., 2001).

While *Fat4* is strongly expressed in the developing retina, we did not observe any major changes to retina morphology or cell composition in *Fat4* mutant mice. The lack of *Fat4^KO^* phenotype in the retina may be due to redundancy of the other three Fat cadherins (Aviles et al., 2022; Aviles et al., 2025; Krol et al., 2016; Sugiyama et al., 2015). *Fat4* may also have functions which we did not explore, such as retinal physiology which may reveal changes in cone/rod function, or the influence of *Fat4* on lens development as signaling from the retina also contributes to proper lens formation (Coomson and Lachke, 2025; Coulombre and Coulombre, 1963).

Previous *in situ* analysis revealed that *Dchs1* transcripts are expressed in the embryonic eye from E9.5-E18.5 (Sugiyama et al., 2015), but its protein localization was unknown. Using an endogenously tagged Dchs1-HA mouse, we detected Dchs1 at the apical surface of the retina neuroepithelium at E14.5. While we observed strong expression of Dchs1 at E14.5 at the apical surface of the neuroblast, we did not observe any overt phenotypes in retinal morphology or cell composition in the P30 mRx-cre; *Dchs1^F/F^*eye. However, given the sequence similarity between *Dchs1* and *Dchs2* and the ability of Dchs2 to bind Fat4, *Dchs2* may be compensating for the loss of *Dchs1*. To explore the compensation hypothesis further experimentation is needed with *Dchs1/2* double mutants.

The outer limiting membrane (OLM) is a belt of adherens junctions that support the connection of photoreceptors to other photoreceptors and to Müller glia cells. Loss of OLM and ONL integrity leads to retinopathies such as Retinitis Pigmentosa and Leber Congenital Amaurosis (Campbell et al., 2007; Heynen et al., 2013; Patel et al., 2026; van de Pavert et al., 2004). *CRB1* mutants, for example, cannot properly form this adherens junction belt resulting in loss of organization of the ONL (van de Pavert et al., 2004). *Dchs2* mutant phenotypes are milder than *CRB1* mutants, but careful analysis of the ONL and OLM revealed changes to the organization of the inner segments and the stacking of the nuclei in the photoreceptors. This suggests that *Dchs2* may play a role in OLM integrity and the ability of photoreceptors to connect to the OLM and to each other. Publicly available single-cell RNA-seq data shows that *Dchs2* is expressed by Müller glia (Aredo et al., 2023). We see a reduction of glutamate synthetase, which marks Müller glia projections, at the OLM in the *Dchs2* mutants. The reduction of Müller glia may contribute to *Dchs2* disorganization, as Müller glia are needed for proper retina lamination (Tworig and Feller, 2021). Alternatively, Dchs2 may directly support the formation or maintenance of the adherens junctions in the OLM.

Taken together, our data show that Fat4, Dchs1 and Dchs2 are dynamically regulated temporally and spatially during embryonic and postnatal eye development; that Fat4 can directly bind to both Dchs1 and Dchs2; and loss of *Fat4*, or *Dchs1* each reduced apical Par3. Most notably we identify a postnatal requirement for *Dchs2*—but not *Fat4* or *Dchs1—*which has a unique role in the columnar organization of photoreceptor nuclei, and the integrity of the outer limiting membrane, defining a *Dchs2* specific, *Fat4*-independent role for this poorly understood Dachsous family member.

## Materials and Methods

### Mice

All mouse procedures were reviewed and approved by the Institutional Animal Care and Use Committee (IACUC) in agreement with animal care guidelines at Washington University- St. Louis. Mouse lines used:

| Mouse Genotype | Identifier |
| --- | --- |
| Fat4<sup>tm1.1Hmc</sup>/B6 | MGI:3846573 |
| Fat4<sup>tm2Hmc</sup> | MGI:3846574 |
| Dchs1<sup>tm1.1Irv</sup>/B6 | MGI:5428667 |
| Dchs1<sup>tm1.2Irv</sup>/B6 | MGI:4867020 |
| Dchs1<sup>em1Rans</sup> | MGI:8280727 |
| Dchs2<sup>tm1.1FHel</sup>/B6 |  |
| Tg(CMV-cre)1Cgn Dchs2<sup>tm1.1FHel</sup> |  |
| C57BL/6J | Strain:000664 |
| Tg(Rax-cre)1Zkoz | MGI:5505625 |

### Generation of *Fat4^Venus^*allele

The mouse model with the tension sensor (mTFP1+mVenus), referred to as *Fat4^Venus^*, fused to the endogenous Fat4 protein was generated by the Genome Engineering & Stem Cell Center (GESC@MGI) and the Mouse Genetics Core at Washington University in St. Louis. Briefly, a gRNA was designed to cleave close to the Fat4 stop codon. Synthetic gRNA with the recognition sequence (PAM): 5’-GCAGATGACGAAGATAATTA(TGG), was validated in mouse N2a cells for efficient cleavage activity. An rAAV donor template was designed to have two 800 bp homology arms flanking the tension sensor coding sequences. The rAAV donor was also validated in N2a cells for targeted integration by junction PCRs. Single-cell mouse embryos were first incubated with the rAAV donor for 5 hours before electroporated with Cas9 protein/sgRNA complex. The surviving embryos were then transferred into pseudo-pregnant females. Live births were screened for positive insertion junctions to identify founders. Founders were then bred with C57Bl/6J mice. F0 pups were screened by toe biopsy genotyping for the tension sensor insert via PCR, see “Genotyping” for primer set.

### Genotyping

Mice were genotyped using the Roche Diagnostic KAPA Mouse Genotyping Kit (Cat. 50-196-5198). DNA was extracted from toe snips of post-natal mice or tail snips of embryonic mice. Genotype was determined by PCR using allele-specific primers.

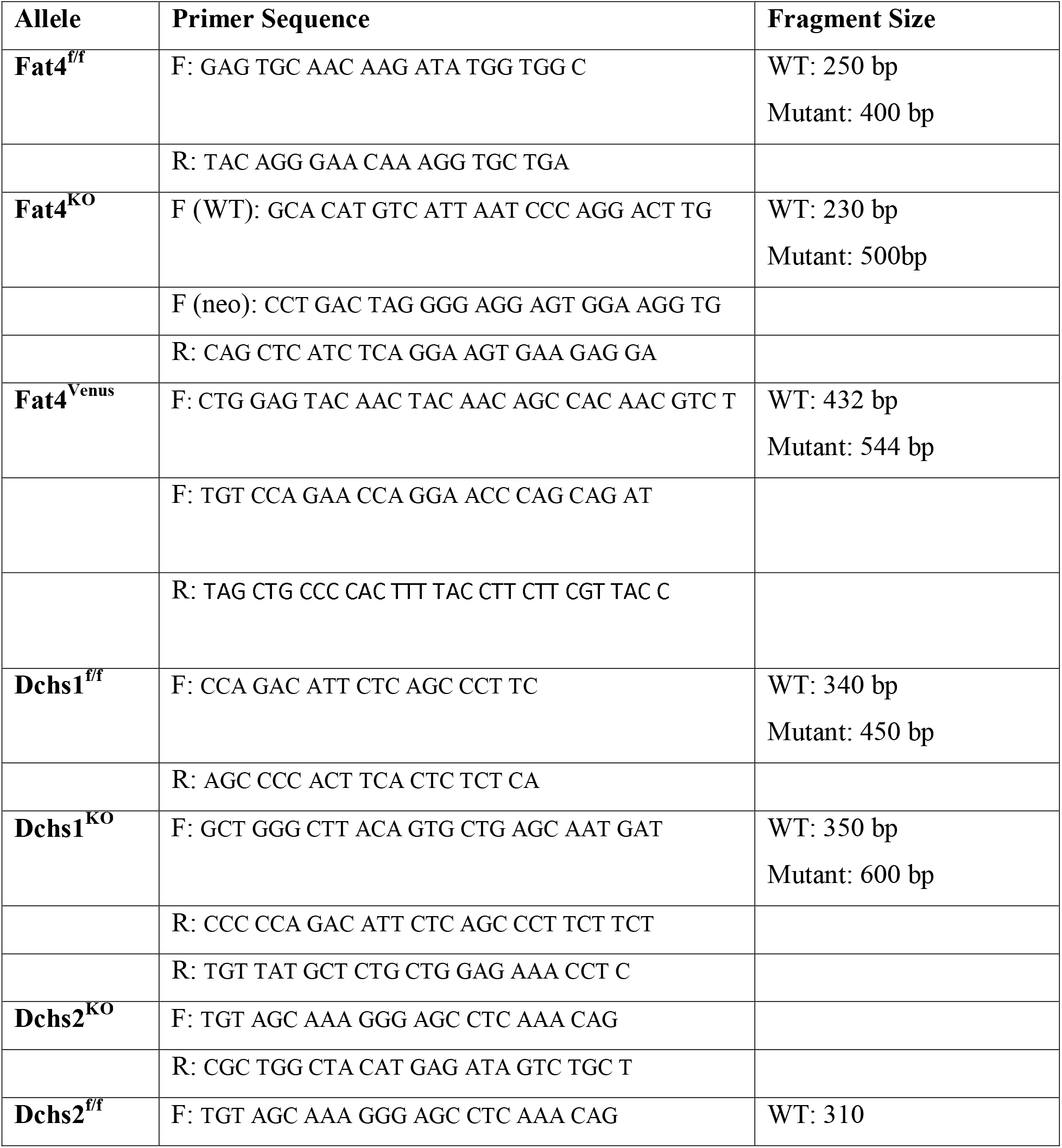
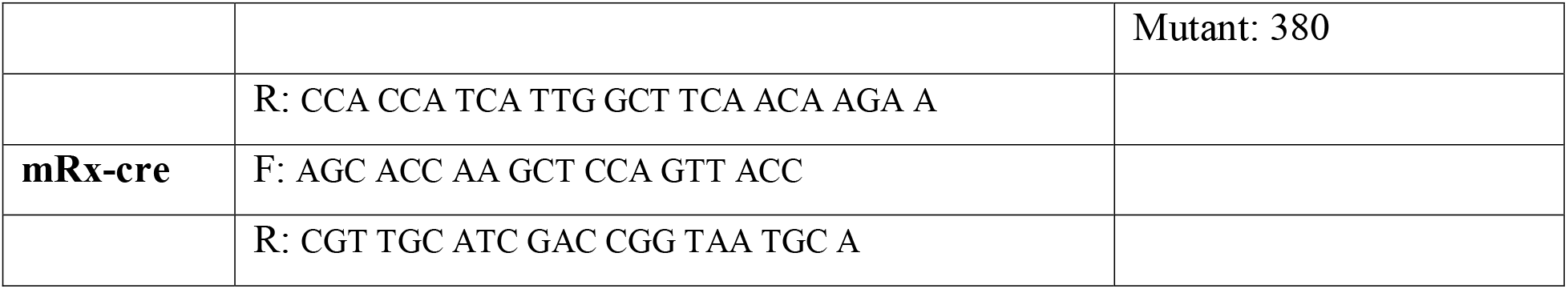

### Western Blot Sample Preparation

For mature retinas (P30 or greater), 1 retina was lysed in 500 uL 2x Laemmli buffer + 2M urea. For retinas P15 or younger 2 retinas were lysed in 500 uL of 2x Laemmli buffer + 2M urea. Samples were bead-beaten using 0.5mm zirconium beads (Next Advance SKU. ZROB05) for 5 minutes at max speed. Samples were dry boiled at 100°C for 10 minutes. Next, samples were centrifuged for 15 minutes, max speed at 4°C. Supernatant was placed in a fresh microcentrifuge tube, flash frozen and stored at −20°C.

### Western Blot Analysis

10-20 uL of lysates were added per lane of NuPage 3-8% Tris-Acetate gels (Thermo Fisher Scientific cat. EA0375BOX) which was determined by normalizing to GAPDH. Samples were run in NuPage Tris-Acetate Running Buffer (Thermo Fisher Scientific cat. LA0041) at 75V for 2 hours. Gels were then wet transferred onto nitrocellulose membranes (Cytiva cat. 10600015) using NuPage Transfer Buffer (Thermo Fisher Scientific cat. NP00061) at 75V for 1.5 hours at 4°C. Membranes were then blocked in 5% milk (RPI cat. M172000) in 1x TBST for 1 hour at room temperature. Primary antibodies were incubated overnight at 4°C. Secondary antibodies were incubated for 1 hour at room temperature. 3, 10 minute washes in 1x TBST were done between primary and secondary antibody incubation and between secondary antibody incubation and visualization. Membranes were visualized via chemiluminescence using SuperSignal West Femto substrate (Thermo Fisher Scientific cat. 1859023) for visualizing Fat4, Dchs1 and Dchs2. SuperSignal West Pico substrate (Thermo Fisher Scientific cat. 34580) was used to visualize GAPDH. See immunohistochemistry antibody dilutions.

### Eye Fixation

For paraformaldehyde (PFA) fixation, 16% PFA (Thermo Fisher cat. 28908) was diluted to 4% in 1x PBS. Once eyes were removed the cornea was pierced with a 22g needle. Eyes were then incubated in 4% PFA for 4 hours at 4 °C on an orbital rocker. Embryos were taken between E14.5 and E15.5, heads were removed and placed in 4% PFA for 24 hours at 4 °C on an orbital rocker. Post fixation, eyes or heads were washed 3x for 10 minutes in 1x PBS. Eyes or heads where then placed in 20% sucrose for 24 hours. Post sucrose incubation eyes or heads were embedded in OCT, frozen in dry ice/ 2-methylbutane, then stored at −80°C till sectioning.

For freeze-substitution, eyes were removed and the cornea punctured with a 22g needle. Eyes were then placed in 3% glacial acetic acid in methanol (M-AA) and placed in a −80 °C freezer for 48 hours. Next eyes were moved to a −20 °C freezer for 4 hours, then left at room temperature for 48 hours. After 48 hours at room temperature, M-AA was replaced with 100 % ethanol. Post fixation eyes were paraffin-embedded and cut to 20 µm thick sections.

### Cell Culture

Cells were cultured in tissue-culture grade flasks with Dulbecco’s Modified Eagle Medium (DMEM) containing 10% fetal bovine serum (FBS) and 1% penicillin-streptomycin. Cells were passaged every three to four days by addition of trypsin-EDTA to dishes and subsequent splitting of cells at a ratio of 1:3

For Par3 co-IP experiments the Flp-InTm T-RExTm Kit was used to generate stable, inducible mammalian expression cell lines by Flp Recombinase-mediated integration. Flp-InTm T-RExTm cell lines generated during this project used the pcDNATm5/FRT/TO expression vector with the gene of interest under control of a tetracycline-regulated, hybrid human cytomegalovirus (CMV)/TetO2 promoter. The vector used also contains the hygromycin resistance gene with an FRT site in the 5’ coding region. Under control of the CMV promoter, the pOG44 plasmid constitutively expresses the Flp recombinase. Plasmids are co-transfected into the Flp-InTm T-RExTm host cell line, at which point the Flp recombinase mediates a homologous recombination event between the FRT sites such that the pcDNATm5/FRT/TO construct is inserted into the genome at the integrated FRT site.

### Plasmids

| Plasmid | Source |
| --- | --- |
| Dchs1-mCherry | AddGene #112855 |
| Fat4-YFP | Gift from David Sprinzak Lab |

### Dchs2-mCherry generation

The Dchs2-mCherry construct was designed using the GenScript gene synthesis design tool and plasmid cloning and prep were done by GenScript. In brief, an mCherry tag was added to the c-terminus of Dchs2 using a GS-linker complex, this was then cloned into pUC57 vector under a CMV promoter. The resulting plasmid was then used for transfection using Lipofectamine 3000.

### Stable cell line generation for Fat4-YFP, Dchs1-mCherry, and Dchs2-mCherry

For the generation of stable cells lines Hek293T cells were transfected using Lipofectamine 3000 (Thermo Fisher cat. L3000015) using the kit provided protocol. Post 2-day incubation of the HEK293T cells with Fat4-YFP (gift from the David Sprinzak Lab), Dchs1-mCherry (Addgene cat. 112855), or Dchs2-mCherry (McNeill lab generated) lipofectamine complex, cells were then treated with selection antibiotic (Fat4-YFP Zeocin 250µg/mL, Dchs1/Dchs2-mCherry Hygromycin 200µg/mL) and maintained in selection media for 10-days. Single cell colonies were generated by limiting dilution in a 96-well plate (2 cells/ml). After a period of two weeks, the plates were screened for positive clones, which were transferred to a new plate for further expansion.

### Cell mixing experiments, immune staining, quantitation and stats

Untransfected (UT) Hek293T cells were mixed at a 50:50 ratio with Fat4-YFP, Dchs1-mCherry, or Dchs2-mCherry expressing cells in a microcentrifuge tube and seeded on glass-bottom, Poly-L-lysine treated plates at a density of 0.05 x 10^6^ cells/well. For cell mixing experiments using UT + Fat4-YPF + Dchs1 or Dchs2-mCherry, cells were mixed at a 1:1:1 ratio. Plates were shaken in a 37°C incubator at 100rpm for 1 hour. Cells were then cultured for 24 hours in standard cell culture conditions. After 24 hours cells were fixed in 4% PFA for 10min, then rinsed twice using 1x PBS. Post fixations cells were blocked for 30min in 10 % donkey sera in 0.5% Triton-X 1x PBS, followed by 1 hour incubation in primary antibody diluted in the blocking buffer, followed by a 30min incubation in secondaries diluted in blocking buffer. Cells were then rinsed twice with 1x PBS. Before imaging, 300µL of fresh 1x PBS was added to the cells.

Protein accumulation of either Fat4-YPF, Dchs1- or Dchs2-mCherry at cell-cell contacts were assed via a line-intensity profile using NIS Elements AR analysis software. A line was drawn from nucleus to nucleus of two contacting cells bisecting the contacting cell membranes. Fluorescent intensity across the 100 contacts counted per condition were averaged and plotted. Statistical analysis was performed using an unpaired t-test using Welch’s correction, with error bars representing SEM. ****= p<0.0001

### Co-Immunoprecipitation

FLAG-Immunoprecipitation was performed 24 to 48 hours after induction Cells were harvested and lysed using the following lysis buffer:

|  |  |
| --- | --- |
| **Lysis Buffer** |  |
| HEPES-NaOH<br>pH=8 | 50mM |
| KCl | 100mM |
| EDTA | 2mM |
| NP-40 | 0.10% |
| Glycerol | 10% |
| ddH2O | to total volume of<br>100mL |
| **Just before use** |  |
| PMSF | 1mM (from 100X) |
| Protease Inhibitor<br>Mix | 500x |

Cells were rinsed once with cold PBS, and cold lysis buffer was added to the cells (0.5 ml/well or 1 ml/10cm plate). Lysates were collected in 1.5 ml microfuge tubes and incubated with shaking for 20 minutes at 4°C. These tubes were then centrifuged at high speed (14,000 rpm) for 20 minutes to pellet the cell debris. As the input control, 40 µl of supernatant from each sample was taken and 10 µl of 5 x SDS buffer was added before boiling for 5 minutes. The remainder of the lysis supernatant was added to 20-30 µl of 50% slurry containing anti-FLAG M2 agarose beads and lysis buffer. (To create the 50% slurry, beads were prewashed 5 times in lysis buffer, then resuspended with the same volume of lysis buffer before use in IP.) The lysates and beads were incubated for 2 hours at 4°C on the nutator. After incubation with beads, lysates were spun down for 10 seconds at low speed (3000 rpm). At this step, a post-IP control sample of 40 µl was taken from each incubated lysate and 10 µl of SDS 5 x buffer was added before boiling for 5 minutes. Remaining lysate was removed and beads were 31 washed 4 times with 1 ml cold lysis buffer, with brief spins at low speed between washes. At the last wash step, all lysis buffer was removed using flat-tip pipette tips. To the dry beads, 30-35 µl of 2 x SDS buffer was added before boiling samples for 5 minutes. Protein extracts were then either stored at −20°C or used in Western blotting experiments

### Immunohistochemistry

PFA fixed eyes or embryonic heads were sectioned to 20 µm thickness. All incubations were done at room temperature. Sections were permeabilized for 1 hour in 0.5% Triton-X 1x PBS. Sections were blocked for 1 hour in 10% donkey sera in 0.5 % Triton-X 1x PBS, sections where then incubated in primary antibody overnight. Next day sections were washed 3 times with 0.5% Triton-X PBS, secondary antibodies were incubated for 2 hours. Sections were then washed 3 times for 5 minutes using 1x PBS. Slides were mounted using Dako Fluorescent Mounting Medium (Millipore Sigma cat. F4680). See table below for antibodies and concentrations used.

| Antibody | Company | Catalog | Host Species | IF Dilution | Western Dilution |
| --- | --- | --- | --- | --- | --- |
| Anti-Dchs1 | Proteintech | 28144-1-AP | Rabbit | N/A | 1:1000 |
| Anti-Dchs2 | McNeill Lab |  | Rabbit | N/A | 1:500 |
| Anti-Fat4 | Millipore-Sigma | HPA052819 | Rabbit | 1:100 | 1:500 |
| Anti-GAPDH | Millipore-Sigma | ABS16 | Rabbit | N/A | 1:10,000 |
| Anti-GFP | Abcam | Ab290 | Rabbit | 1:1000 | 1:10,000 |
| Anti-mCherry | Abcam | ab167453 | Rabbit | 1:500 | 1:1000 |
| Anti-mCherry | Abcam | ab205402 | Chicken | 1:500 | N/A |
| Anti-MCT1 | Gift from Nancy Phillip Lab |  | Rabbit | 1:250 | N/A |
| Anti-P Cadherin | R&D Systems | AF761 | Goat | 1:500 | N/A |
| Anti-Pard3 | Millipore-Sigma | HPA030443 | Rabbit | 1:200 | N/A |
| Calbindin D28K | Synaptic Systems | 214 004 | Guinea pig | 1:500 | N/A |
| Cone Arrestin | Millipore-Sigma | Ab15282 | Rabbit | 1:500 | N/A |
| Glutamine Synthetase | Millipore-Sigma | GS2781 | Rabbit | 1:500 | N/A |
| PKCα | Santa Cruz | SC-8393 | Mouse | 1:500 | N/A |
| Rhodopsin | Santa Cruz | SC-57432 | Mouse | 1:500 | N/A |
| Yap/Taz (D24E4) | Cell signaling | 8418S | Rabbit | 1:100 | N/A |
| ZO-1 | Invitrogen | 61-7300 | Rabbit | 1:300 | N/A |

| Secondary/Dye | Company | Catalog | IF Dilution | Western Dilution |
| --- | --- | --- | --- | --- |
| Donkey anti-rabbit FAB | Jackson ImmunoResearch | 711-547-003 | 1:5,000 | N/A |
| Dk anti-mouse 594 | ThermoFisher | A-21203 | 1:500 | N/A |
| Dk anti-goat 594 | ThermoFisher | A-11058 | 1:500 | N/A |
| Dk anti-goat 647 | ThermoFisher | A-21447 | 1:500 | N/A |
| Dk anti-Chicken 594 | ThermoFisher | A78591 | 1:500 | N/A |
| DAPI | ThermoFisher | 62248 | 1:1,000 | N/A |
| Phalloidin 594 | ThermoFisher | A12381 | 1:500 | N/A |
| Phalloidin 633 | ThermoFisher | A22284 | 1:500 | N/A |
| Anti-Rabbit HRP | Proteintech | SA00001-9 | N/A | 1:5000 |

### Morphology and Cell Count Analysis

For retina thickness, 20x images of hematoxylin and eosin stained sections were stitched together, and the layers of the retina were measured using ImageJ software. The thickness of the retinal layers were measured at regular distances from the optic nerve. For cell type quantification, 3 animals were collected for each genotype and 3 sections per eye were stained and imaged. Statistical analyses were performed using two-way ANOVA with multiple comparisons. All statistical analysis was performed using GraphPad Prism 10.

### Software and Statistical Analysis

| Product | Source | Identifier |
| --- | --- | --- |
| ImageJ | ImageJ (v1.54p) | https://imagej.net/ij/download.html |
| Prism | GraphPad (v10) | https://www.graphpad.com/ |
| NIS Elements AR Analysis | Nikon (v5.30.05) | https://www.microscope.healthcare.nikon.com/products/software/nis-elements |
| Photoshop/ Illustrator | Adobe Creative Cloud (v6.6.0.611) | https://www.adobe.com/home |
| EndNote | EndNote (v21) | https://endnote.com/ |

Quantifications are reported as mean ± SD, all figures were generated using Prism v.10. Statistical tests are indicated within figure legends. n values represent the number of biological replicates, independent animals. Animal numbers, cell counts, and statistical methods can be found in figure legends.

## References

Abuderman, A.A., Harb, O.A., Gertallah, L.M., 2020. Prognostic and clinic-pathological significances of HOXB8, ILK and FAT4 expression in colorectal cancer. Contemp Oncol (Pozn) 24, 183–192.

Adler, R., Belecky-Adams, T.L., 2002. The role of bone morphogenetic proteins in the differentiation of the ventral optic cup. Development 129, 3161–3171.

Ambegaonkar, A.A., Pan, G., Mani, M., Feng, Y., Irvine, K.D., 2012. Propagation of Dachsous-Fat planar cell polarity. Curr Biol 22, 1302–1308.

Angst, B.D., Marcozzi, C., Magee, A.I., 2001. The cadherin superfamily: diversity in form and function. J Cell Sci 114, 629–641.

Aredo, B., Kumar, A., Chen, B., Xing, C., Ufret-Vincenty, R.L., 2023. Single Cell RNA Sequencing Analysis of Mouse Retina Identifies a Subpopulation of Muller Glia Involved in Retinal Recovery From Injury in the FCD-LIRD Model. Invest Ophthalmol Vis Sci 64, 2.

Aviles, E.C., Krol, A., Henle, S.J., Burroughs-Garcia, J., Deans, M.R., Goodrich, L.V., 2022. Fat3 acts through independent cytoskeletal effectors to coordinate asymmetric cell behaviors during polarized circuit assembly. Cell Rep 38, 110307.

Aviles, E.C., Wang, S.K., Patel, S., Cordero, S., Shi, S., Lin, L., Kefalov, V.J., Goodrich, L.V., Cepko, C.L., Xue, Y., 2025. ERG responses to high-frequency flickers require FAT3 signaling in mouse retinal bipolar cells. J Gen Physiol 157.

Badouel, C., Zander, M.A., Liscio, N., Bagherie-Lachidan, M., Sopko, R., Coyaud, E., Raught, B., Miller, F.D., McNeill, H., 2015. Fat1 interacts with Fat4 to regulate neural tube closure, neural progenitor proliferation and apical constriction during mouse brain development. Development 142, 2781–2791.

Baena-Lopez, L.A., Rodriguez, I., Baonza, A., 2008. The tumor suppressor genes dachsous and fat modulate different signalling pathways by regulating dally and dally-like. Proc Natl Acad Sci U S A 105, 9645–9650.

Bagherie-Lachidan, M., Reginensi, A., Pan, Q., Zaveri, H.P., Scott, D.A., Blencowe, B.J., Helmbacher, F., McNeill, H., 2015. Stromal Fat4 acts non-autonomously with Dchs1/2 to restrict the nephron progenitor pool. Development 142, 2564–2573.

Bennett, F.C., Harvey, K.F., 2006. Fat cadherin modulates organ size in Drosophila via the Salvador/Warts/Hippo signaling pathway. Curr Biol 16, 2101–2110.

Bosch, J.A., Sumabat, T.M., Hafezi, Y., Pellock, B.J., Gandhi, K.D., Hariharan, I.K., 2014. The Drosophila F-box protein Fbxl7 binds to the protocadherin fat and regulates Dachs localization and Hippo signaling. Elife 3, e03383.

Bosveld, F., Bonnet, I., Guirao, B., Tlili, S., Wang, Z., Petitalot, A., Marchand, R., Bardet, P.L., Marcq, P., Graner, F., Bellaiche, Y., 2012. Mechanical control of morphogenesis by Fat/Dachsous/Four-jointed planar cell polarity pathway. Science 336, 724–727.

Brittle, A., Warrington, S.J., Strutt, H., Manning, E., Tan, S.E., Strutt, D., 2022. Distinct mechanisms of planar polarization by the core and Fat-Dachsous planar polarity pathways in the Drosophila wing. Cell Rep 40, 111419.

Bryant, P.J., Huettner, B., Held, L.I., Jr., Ryerse, J., Szidonya, J., 1988. Mutations at the fat locus interfere with cell proliferation control and epithelial morphogenesis in Drosophila. Dev Biol 129, 541–554.

Byerly, K., Wolfe, C., Parris, H., Griggs, C., Wilson, E., Huff, M., Griggs, M., Morningstar, J., Guo, L., Tang, F., Guz, J., Petrucci, T., Phookan, R., Loizzi, B., Gensemer, C., Norris, R.A., 2025. Dynamic Expression and Functional Implications of the Cell Polarity Gene, Dchs1, During Cardiac Development. Cells 14.

Cai, J., Feng, D., Hu, L., Chen, H., Yang, G., Cai, Q., Gao, C., Wei, D., 2015. FAT4 functions as a tumour suppressor in gastric cancer by modulating Wnt/beta-catenin signalling. Br J Cancer 113, 1720–1729.

Campbell, M., Humphries, M., Kenna, P., Humphries, P., Brankin, B., 2007. Altered expression and interaction of adherens junction proteins in the developing OLM of the Rho(-/-) mouse. Exp Eye Res 85, 714–720.

Cappello, S., Gray, M.J., Badouel, C., Lange, S., Einsiedler, M., Srour, M., Chitayat, D., Hamdan, F.F., Jenkins, Z.A., Morgan, T., Preitner, N., Uster, T., Thomas, J., Shannon, P., Morrison, V., Di Donato, N., Van Maldergem, L., Neuhann, T., Newbury-Ecob, R., Swinkells, M., Terhal, P., Wilson, L.C., Zwijnenburg, P.J., Sutherland-Smith, A.J., Black, M.A., Markie, D., Michaud, J.L., Simpson, M.A., Mansour, S., McNeill, H., Gotz, M., Robertson, S.P., 2013. Mutations in genes encoding the cadherin receptor-ligand pair DCHS1 and FAT4 disrupt cerebral cortical development. Nat Genet 45, 1300–1308.

Castelvecchi, G., Li, L., Shin, J., Li-Villarreal, N., Roszko, I., Gontarz, P., Li, T., Zhang, B., Sepich, D.S., Solnica-Krezel, L., 2026. Regulation of the yolk microtubule and actin cytoskeleton by Dachsous cadherins during zebrafish epiboly. Dev Biol 537, 71–86.

Che, X., Jian, F., Jia, N., Zheng, Y., Jiang, Y., Feng, W., 2019. FAT4-USP51 complex regulates the proliferation and invasion of endometrial cancer via Hippo pathway. Am J Transl Res 11, 2784–2800.

Cho, E., Feng, Y., Rauskolb, C., Maitra, S., Fehon, R., Irvine, K.D., 2006. Delineation of a Fat tumor suppressor pathway. Nat Genet 38, 1142–1150.

Clark, H.F., Brentrup, D., Schneitz, K., Bieber, A., Goodman, C., Noll, M., 1995. Dachsous encodes a member of the cadherin superfamily that controls imaginal disc morphogenesis in Drosophila. Genes Dev 9, 1530–1542.

Coomson, S.Y., Lachke, S.A., 2025. Bringing signaling complexity into focus. Elife 14.

Coulombre, J.L., Coulombre, A.J., 1963. Lens Development: Fiber Elongation and Lens Orientation. Science 142, 1489–1490.

Crespo-Enriquez, I., Hodgson, T., Zakaria, S., Cadoni, E., Shah, M., Allen, S., Al-Khishali, A., Mao, Y., Yiu, A., Petzold, J., Villagomez-Olea, G., Pitsillides, A.A., Irvine, K.D., Francis-West, P., 2019. Dchs1-Fat4 regulation of osteogenic differentiation in mouse. Development 146.

Du, J., Ji, J., Gao, Y., Xu, L., Xu, J., Zhu, C., Gu, H., Jiang, J., Li, H., Ma, H., Hu, Z., Jin, G., Guo, W., Chen, X., Shen, H., 2013. Nonsynonymous polymorphisms in FAT4 gene are associated with the risk of esophageal cancer in an Eastern Chinese population. Int J Cancer 133, 357–361.

Ebnet, K., Suzuki, A., Horikoshi, Y., Hirose, T., Meyer Zu Brickwedde, M.K., Ohno, S., Vestweber, D., 2001. The cell polarity protein ASIP/PAR-3 directly associates with junctional adhesion molecule (JAM). EMBO J 20, 3738–3748.

Fulford, A., Enderle, L., Rusch, J., Hodzic, D., Holder, M.V., Earl, A., Hyunseo Oh, R., Tapon, N., McNeill, H., 2023. Expanded directly binds conserved regios of Fat to restrain growth via the Hippo pathway J Cell Biol p. in press.

Fulford, A.D., McNeill, H., 2020. Fat/Dachsous family cadherins in cell and tissue organisation. Curr Opin Cell Biol 62, 96–103.

Heynen, S.R., Meneau, I., Caprara, C., Samardzija, M., Imsand, C., Levine, E.M., Grimm, C., 2013. CDC42 is required for tissue lamination and cell survival in the mouse retina. PLoS One 8, e53806.

Hou, L., Chen, M., Zhao, X., Li, J., Deng, S., Hu, J., Yang, H., Jiang, J., 2016. FAT4 functions as a tumor suppressor in triple-negative breast cancer. Tumour Biol.

Hulpiau, P., van Roy, F., 2009. Molecular evolution of the cadherin superfamily. Int J Biochem Cell Biol 41, 349–369.

Ishiuchi, T., Misaki, K., Yonemura, S., Takeichi, M., Tanoue, T., 2009. Mammalian Fat and Dachsous cadherins regulate apical membrane organization in the embryonic cerebral cortex. J Cell Biol 185, 959–967.

Izumi, Y., Hirose, T., Tamai, Y., Hirai, S., Nagashima, Y., Fujimoto, T., Tabuse, Y., Kemphues, K.J., Ohno, S., 1998. An atypical PKC directly associates and colocalizes at the epithelial tight junction with ASIP, a mammalian homologue of Caenorhabditis elegans polarity protein PAR-3. J Cell Biol 143, 95–106.

Jiang, X., Liu, Z., Xia, Y., Luo, J., Xu, J., He, X., Tao, H., 2018. Low FAT4 expression is associated with a poor prognosis in gastric cancer patients. Oncotarget 9, 5137–5154.

Joberty, G., Petersen, C., Gao, L., Macara, I.G., 2000. The cell-polarity protein Par6 links Par3 and atypical protein kinase C to Cdc42. Nat Cell Biol 2, 531–539.

Kanda, A., Noda, K., Yuki, K., Ozawa, Y., Furukawa, T., Ichihara, A., Ishida, S., 2013. Atp6ap2/(pro)renin receptor interacts with Par3 as a cell polarity determinant required for laminar formation during retinal development in mice. J Neurosci 33, 19341–19351.

Klimova, L., Lachova, J., Machon, O., Sedlacek, R., Kozmik, Z., 2013. Generation of mRx-Cre transgenic mouse line for efficient conditional gene deletion in early retinal progenitors. PLoS One 8, e63029.

Krol, A., Henle, S.J., Goodrich, L.V., 2016. Fat3 and Ena/VASP proteins influence the emergence of asymmetric cell morphology in the developing retina. Development 143, 2172–2182.

Kuta, A., Mao, Y., Martin, T., Ferreira de Sousa, C., Whiting, D., Zakaria, S., Crespo-Enriquez, I., Evans, P., Balczerski, B., Mankoo, B., Irvine, K.D., Francis-West, P.H., 2016. Fat4-Dchs1 signalling controls cell proliferation in developing vertebrae. Development 143, 2367–2375.

Li, Q., Zhou, X., Fang, Z., Pan, Z., Zhou, H., 2020. Up-regulation of FAT4 enhances the chemosensitivity of colorectal cancer cells treated by 5-FU. Transl Cancer Res 9, 309–322.

Li, S.Y., Wang, H., Mai, H.F., Li, G.F., Chen, S.J., Li, G.S., Liang, B.C., 2019. Down-regulated long non-coding RNA RNAZFHX4-AS1 suppresses invasion and migration of breast cancer cells via FAT4-dependent Hippo signaling pathway. Cancer Gene Ther 26, 374–387.

Lin, D., Edwards, A.S., Fawcett, J.P., Mbamalu, G., Scott, J.D., Pawson, T., 2000. A mammalian PAR-3-PAR-6 complex implicated in Cdc42/Rac1 and aPKC signalling and cell polarity. Nat Cell Biol 2, 540–547.

Lodge, E.J., Xekouki, P., Silva, T.S., Kochi, C., Longui, C.A., Faucz, F.R., Santambrogio, A., Mills, J.L., Pankratz, N., Lane, J., Sosnowska, D., Hodgson, T., Patist, A.L., Francis-West, P., Helmbacher, F., Stratakis, C., Andoniadou, C.L., 2020. Requirement of FAT and DCHS protocadherins during hypothalamic-pituitary development. JCI Insight 5.

Loza, O., Heemskerk, I., Gordon-Bar, N., Amir-Zilberstein, L., Jung, Y., Sprinzak, D., 2017. A synthetic planar cell polarity system reveals localized feedback on Fat4-Ds1 complexes. Elife 6.

Ma, D., Yang, C.H., McNeill, H., Simon, M.A., Axelrod, J.D., 2003. Fidelity in planar cell polarity signalling. Nature 421, 543–547.

Ma, L., Cui, J., Xi, H., Bian, S., Wei, B., Chen, L., 2016. Fat4 suppression induces Yap translocation accounting for the promoted proliferation and migration of gastric cancer cells. Cancer Biol Ther 17, 36–47.

Malgundkar, S.H., Burney, I., Al Moundhri, M., Al Kalbani, M., Lakhtakia, R., Okamoto, A., Tamimi, Y., 2020. FAT4 silencing promotes epithelial-to-mesenchymal transition and invasion via regulation of YAP and beta-catenin activity in ovarian cancer. BMC Cancer 20, 374.

Mao, W., Zhou, J., Hu, J., Zhao, K., Fu, Z., Wang, J., Mao, K., 2022. A pan-cancer analysis of FAT atypical cadherin 4 (FAT4) in human tumors. Front Public Health 10, 969070.

Mao, Y., Kuta, A., Crespo-Enriquez, I., Whiting, D., Martin, T., Mulvaney, J., Irvine, K.D., Francis-West, P., 2016. Dchs1-Fat4 regulation of polarized cell behaviours during skeletal morphogenesis. Nat Commun 7, 11469.

Mao, Y., Mulvaney, J., Zakaria, S., Yu, T., Morgan, K.M., Allen, S., Basson, M.A., Francis-West, P., Irvine, K.D., 2011. Characterization of a Dchs1 mutant mouse reveals requirements for Dchs1-Fat4 signaling during mammalian development. Development 138, 947–957.

Matakatsu, H., Blair, S.S., 2004. Interactions between Fat and Dachsous and the regulation of planar cell polarity in the Drosophila wing. Development 131, 3785–3794.

Matakatsu, H., Blair, S.S., 2006. Separating the adhesive and signaling functions of the Fat and Dachsous protocadherins. Development 133, 2315–2324.

Matakatsu, H., Blair, S.S., 2012. Separating planar cell polarity and Hippo pathway activities of the protocadherins Fat and Dachsous. Development 139, 1498–1508.

Matis, M., Axelrod, J.D., 2013. Regulation of PCP by the Fat signaling pathway. Genes Dev 27, 2207–2220.

Medina, E., Easa, Y., Lester, D.K., Lau, E.K., Sprinzak, D., Luca, V.C., 2023. Structure of the planar cell polarity cadherins Fat4 and Dachsous1. Nat Commun 14, 891.

Molday, L.L., Wu, W.W., Molday, R.S., 2007. Retinoschisin (RS1), the protein encoded by the X-linked retinoschisis gene, is anchored to the surface of retinal photoreceptor and bipolar cells through its interactions with a Na/K ATPase-SARM1 complex. J Biol Chem 282, 32792–32801.

Moon, K.H., Kim, H.T., Lee, D., Rao, M.B., Levine, E.M., Lim, D.S., Kim, J.W., 2018. Differential Expression of NF2 in Neuroepithelial Compartments Is Necessary for Mammalian Eye Development. Dev Cell 44, 13–28 e13.

Nagai-Tamai, Y., Mizuno, K., Hirose, T., Suzuki, A., Ohno, S., 2002. Regulated protein-protein interaction between aPKC and PAR-3 plays an essential role in the polarization of epithelial cells. Genes Cells 7, 1161–1171.

Pan, M., Chen, Q., Lu, Y., Wei, F., Chen, C., Tang, G., Huang, H., 2020. MiR-106b-5p regulates the migration and invasion of colorectal cancer cells by targeting FAT4. Biosci Rep 40.

Patel, M.K., Piedade, W., Famulski, J.K., 2026. Cdhr1a and pcdh15b link photoreceptor outer segments with inner segment calyceal processes revealing a potential mechanism for cone-rod dystrophy. bioRxiv.

Ragni, C.V., Diguet, N., Le Garrec, J.F., Novotova, M., Resende, T.P., Pop, S., Charon, N., Guillemot, L., Kitasato, L., Badouel, C., Dufour, A., Olivo-Marin, J.C., Trouve, A., McNeill, H., Meilhac, S.M., 2017. Amotl1 mediates sequestration of the Hippo effector Yap1 downstream of Fat4 to restrict heart growth. Nat Commun 8, 14582.

Rock, R., Schrauth, S., Gessler, M., 2005. Expression of mouse dchs1, fjx1, and fat-j suggests conservation of the planar cell polarity pathway identified in Drosophila. Dev Dyn 234, 747–755.

Saburi, S., Hester, I., Fischer, E., Pontoglio, M., Eremina, V., Gessler, M., Quaggin, S.E., Harrison, R., Mount, R., McNeill, H., 2008. Loss of Fat4 disrupts PCP signaling and oriented cell division and leads to cystic kidney disease. Nat Genet 40, 1010–1015.

Sai, X., Ikawa, Y., Nishimura, H., Mizuno, K., Kajikawa, E., Katoh, T.A., Kimura, T., Shiratori, H., Takaoka, K., Hamada, H., Minegishi, K., 2022. Planar cell polarity-dependent asymmetric organization of microtubules for polarized positioning of the basal body in node cells. Development 149.

Sharma, P., McNeill, H., 2013. Regulation of long-range planar cell polarity by Fat-Dachsous signaling. Development 140, 3869–3881.

Silva, E., Tsatskis, Y., Gardano, L., Tapon, N., McNeill, H., 2006. The tumor-suppressor gene fat controls tissue growth upstream of expanded in the hippo signaling pathway. Curr Biol 16, 2081–2089.

Sing, A., Tsatskis, Y., Fabian, L., Hester, I., Rosenfeld, R., Serricchio, M., Yau, N., Bietenhader, M., Shanbhag, R., Jurisicova, A., Brill, J.A., McQuibban, G.A., McNeill, H., 2014. The atypical cadherin fat directly regulates mitochondrial function and metabolic state. Cell 158, 1293–1308.

Strutt, H., Meshram, D., Manning, E., Madathil, A.C.K., Strutt, D., 2024. Fat-Dachsous planar polarity function requires two distinct heterophilic cadherin-cadherin binding interactions. Cell Rep 43, 114722.

Sugiyama, Y., Lovicu, F.J., McAvoy, J.W., 2011. Planar cell polarity in the mammalian eye lens. Organogenesis 7, 191–201.

Sugiyama, Y., Shelley, E.J., Badouel, C., McNeill, H., McAvoy, J.W., 2015. Atypical Cadherin Fat1 Is Required for Lens Epithelial Cell Polarity and Proliferation but Not for Fiber Differentiation. Invest Ophthalmol Vis Sci 56, 4099–4107.

Sugiyama, Y., Stump, R.J., Nguyen, A., Wen, L., Chen, Y., Wang, Y., Murdoch, J.N., Lovicu, F.J., McAvoy, J.W., 2010. Secreted frizzled-related protein disrupts PCP in eye lens fiber cells that have polarised primary cilia. Dev Biol 338, 193–201.

Sun, H., Zhou, H., Zhang, Y., Chen, J., Han, X., Huang, D., Ren, X., Jia, Y., Fan, Q., Tian, W., Zhao, Y., 2018. Aberrant methylation of FAT4 and SOX11 in peripheral blood leukocytes and their association with gastric cancer risk. J Cancer 9, 2275–2283.

Tanoue, T., Takeichi, M., 2005. New insights into Fat cadherins. J Cell Sci 118, 2347–2353.

Tworig, J.M., Feller, M.B., 2021. Muller Glia in Retinal Development: From Specification to Circuit Integration. Front Neural Circuits 15, 815923.

van de Pavert, S.A., Kantardzhieva, A., Malysheva, A., Meuleman, J., Versteeg, I., Levelt, C., Klooster, J., Geiger, S., Seeliger, M.W., Rashbass, P., Le Bivic, A., Wijnholds, J., 2004. Crumbs homologue 1 is required for maintenance of photoreceptor cell polarization and adhesion during light exposure. J Cell Sci 117, 4169–4177.

Wei, R., Xiao, Y., Song, Y., Yuan, H., Luo, J., Xu, W., 2019. FAT4 regulates the EMT and autophagy in colorectal cancer cells in part via the PI3K-AKT signaling axis. J Exp Clin Cancer Res 38, 112.

Willecke, M., Hamaratoglu, F., Kango-Singh, M., Udan, R., Chen, C.L., Tao, C., Zhang, X., Halder, G., 2006. The fat cadherin acts through the hippo tumor-suppressor pathway to regulate tissue size. Curr Biol 16, 2090–2100.

Willecke, M., Hamaratoglu, F., Sansores-Garcia, L., Tao, C., Halder, G., 2008. Boundaries of Dachsous Cadherin activity modulate the Hippo signaling pathway to induce cell proliferation. Proc Natl Acad Sci U S A 105, 14897–14902.

Zakaria, S., Mao, Y., Kuta, A., de Sousa, C.F., Gaufo, G.O., McNeill, H., Hindges, R., Guthrie, S., Irvine, K.D., Francis-West, P.H., 2014. Regulation of neuronal migration by Dchs1-Fat4 planar cell polarity. Curr Biol 24, 1620–1627.

Zhang, H., Bagherie-Lachidan, M., Badouel, C., Enderle, L., Peidis, P., Bremner, R., Kuure, S., Jain, S., McNeill, H., 2019. FAT4 Fine-Tunes Kidney Development by Regulating RET Signaling. Dev Cell 48, 780–792 e784.

Zhao, X., Yang, C.H., Simon, M.A., 2013. The Drosophila Cadherin Fat regulates tissue size and planar cell polarity through different domains. PLoS One 8, e62998.

