## Supplemental Figures for "Dchs2, a novel Fat4 ligand, is required for photoreceptor organization and outer limiting membrane integrity in the mouse retina"

Supplement 1

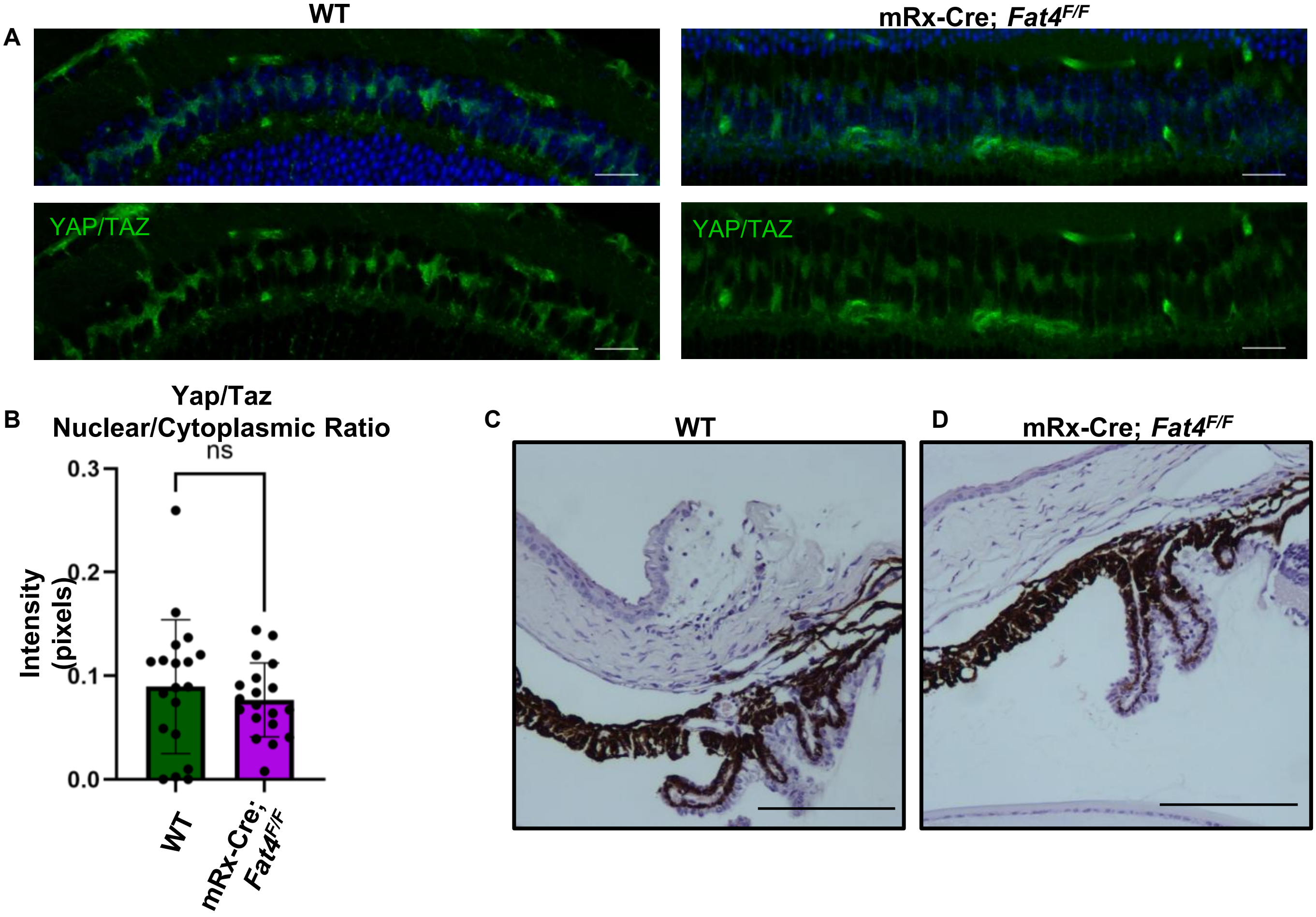


**Supplement 1. Loss of *Fat4* does not affect Hippo pathway readouts**
**A.** WT and Rx-Cre; *Fat4*$^{F/F}$ retinas stained for YAP/TAZ (green) and nuclei (blue) Scale bars= 20µm. **B.** Quantitation of the Yap/Taz cytoplasmic to nucleus ratio shows nonsignificant difference between control and mutant. **C.** WT 20x image of the ciliary body. Rx-Cre; *Fat4*$^{F/F}$ 20x image of the ciliary body. There was no changes to growth of the ciliary in the loss of *Fat4* condition. Scale bars= 50µm

**Supplement 2**

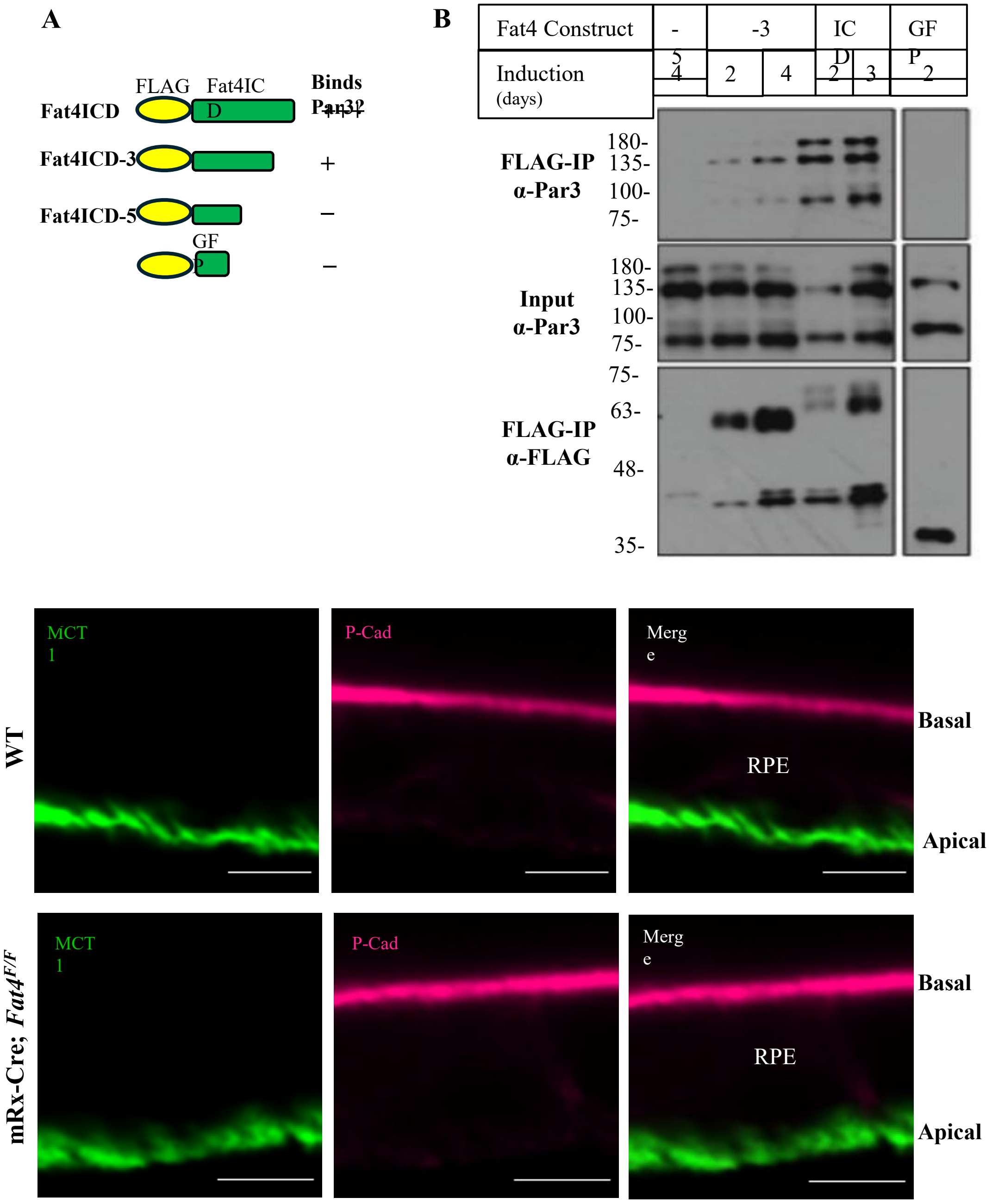


**Supplement 2. Fat4 binds Par3 but loss of Fat4 does not affect apico-basal polarity in the RPE A.** Schematic of FLAG-Fat4 ICD (intracellular domain) constructs and if they bind PAR3. **B.** Western blot of FLAG-FAT4 ICD constructs blotting for Par3, FLAG or GFP. Full-length FLAG-Fat4-ICD bound Par3 the strongest (+++), FLAG-Fat4-ICD-3 weakly bound Par3 (+), and FLAG-Fat4-ICD-5 did not bind Par3 (-). **C.** WT and Rx-Cre; *Fat4*$^{F/F}$ P30 RPE labeled with the basal marker P-cadherin (P-Cad) and the apical marker MCT1 showing no changes in RPE apico-basal localization when *Fat4* is lost.

**Supplement 3**

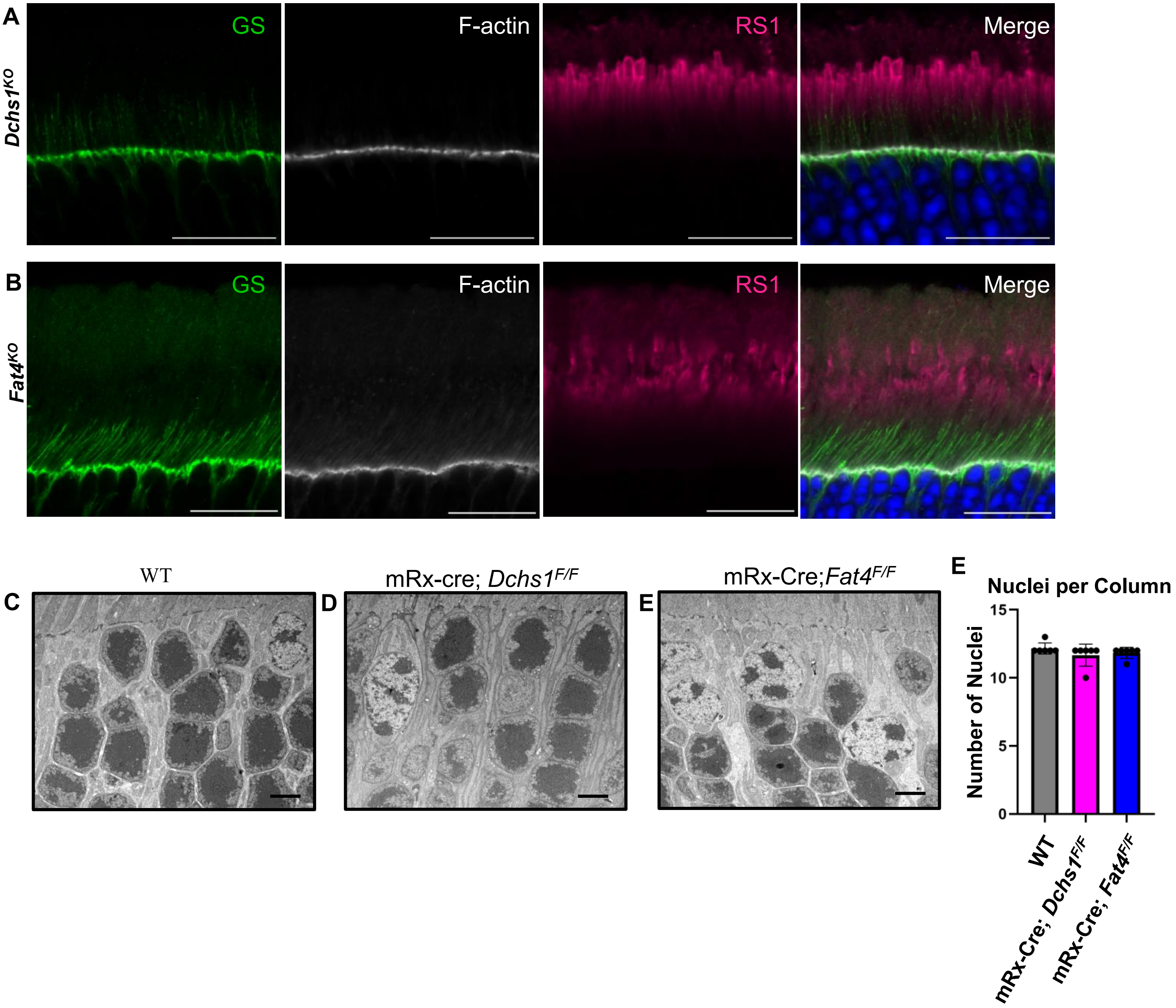


**Supplement 3. Loss of *Fat4* or *Dchs1* does not affect inner segments, OLM integrity or photoreceptor organization**

**A & B.** *Dchs1*$^{KO}$ (**A.**) or *Fat4*$^{KO}$ (**B.**) labeled for glutamine synthetase in green (Müller Glia projections), F-actin in white, and RS1 in pink (inner segments). **C-E** WT (**C**) mRx-cre; *Dchs1*$^{F/F}$ (**D**) or mRx-Cre; *Fat4*$^{F/F}$ (**E**) showing no change in nuclear stacking of the photoreceptors when either *Fat4* or *Dchs1* is lost in the P30 eye. **F.** Quantitation of number of nuclei per column in WT, *Fat4* and *Dchs1* mutant animals
